# TET1 attenuates human pluripotency-related heterogeneities and enhances differentiation capabilities

**DOI:** 10.1101/2022.04.29.489839

**Authors:** Makoto Motono, Keiko Hiraki-Kamon, Masayoshi Kamon, Hidenori Kiyosawa, Yoichi Kondo, Hidemasa Kato

## Abstract

Human induced pluripotent stem cells (hiPSCs) offer transformative potential for developmental research and therapeutic discovery; however, their variable characteristics are a significant barrier to their broader application. We reprogrammed hiPSCs by adding TET1, a DNA-demethylating dioxygenase, to produce TET1-iPSCs (T-iPSCs) with enhanced epithelialization and differentiation capabilities. A default differentiation survey—designed and implemented in our hiPSC-production pipeline—revealed considerable variation in differentiation capacity across qualitatively pluripotent iPSC clones. By comparing high-differentiation T-iPSCs and conventional iPSCs (C-iPSCs) at the single-cell level, we identified a subpopulation of C-iPSCs in the G0/G1 phase (*CDKN2A*-positive) exhibiting low *TET1* expression and an elevated extraembryonic mesenchymal gene signature. The extraembryonic mesenchymal gene signature is common to iPSCs and quantitatively correlates to default differentiation results. Deviation of C-iPSCs from the cell-cycling embryonic epithelial gene profile was accompanied by DNA hypermethylation concomitant with *TET1* insufficiency and the accompanying derepression of poised and Polycomb-repressed enhancers, leading to the derepression of senescence-associated and extraembryonic genes. TET1-facilitated reprogramming can ameliorate these deviations. This study provides a standardized hiPSC- production pipeline and discovers an aspect of TET1 for establishing human pluripotency by remedying the inherent heterogeneities resulting from C-iPSC reprogramming.

## Introduction

Human pluripotent stem cells (hPSCs) exhibit variable capabilities for differentiation, which is more pronounced in human induced PSCs (hiPSCs) (Koyanagi-Aoi et al. 2013; Ohnuki et al. 2014) than in human embryonic stem cells (hESCs) (Bock et al. 2011). Efforts to optimize human reprogramming have focused on enhancing the efficiency of iPSC colony derivation (Esteban et al. 2010; Wang et al. 2017); however, strategies to improve differentiation capacities are limited. In mice, Tet1, a DNA-demethylating dioxygenase, increased the differentiation potential of iPSCs by demethylating enhancers (Bartoccetti et al. 2020). Mesenchymal-to-epithelial transition is an essential step of reprogramming (Schiebinger et al. 2019), in which Tet1 plays a crucial role (Hu et al. 2014). Compared to that in mice, in humans, the onset of *TET1* expression during reprogramming is more delayed, which hinders mesenchymal-to-epithelial transition and leads to epigenetic aberrations (Cacchiarelli et al. 2015; Bartoccetti et al. 2020). Therefore, the Yamanaka factors alone may be insufficient to demethylate the genome of human mesenchymal cells and attain genuine pluripotency.

We propose a novel hiPSC-production pipeline that adds TET1 to the Yamanaka factors, yielding TET1-iPSCs (T-iPSCs). This pipeline integrates differentiation tests of individual clones, selecting hiPSCs with enhanced differentiation capabilities. Assessment of differentiation capabilities and the corresponding systematic transcriptomics revealed inter- and intra-clonal cell heterogeneities incidental to the differentiation defects of conventional hiPSCs (C-iPSCs). Analyses of the DNA methylome and histone modification support the hypothesis that the dysregulation of this gene is inherent to conventional reprogramming from somatic cells.

## Results

### TET1-aided human cell reprogramming: T-iPSCs

During hiPSC reprogramming, the establishment of pluripotent DNA methylation patterns is delayed compared with that in mice (Cacchiarelli et al. 2015). By reanalyzing single-cell transcriptome data from human reprogramming (Liu et al. 2020), we found that *TET1* expression was selectively pronounced during a successful reprogramming, followed by successive *DNMT3A* and *DNMT3B* upregulation (Supplemental Fig. S1A,B). Given the known pleiotropic roles of TET1 in reprogramming and pluripotency (Verma et al. 2018; Bartoccetti et al. 2020; Cheng et al. 2022), we generated hiPSCs using episomal vectors carrying conventional reprogramming factors alone (C-iPSCs) (Okita et al. 2011) and with *TET1* addition (T-iPSCs) (Fig. 1A). We cultured hiPSCs in an epiblast stem cell medium (Brons et al. 2007). This enabled single-cell passage and improved morphological characterization (Fig. 1B). During colony formation, most emerging T-iPSCs organized into a simple columnar epithelium. These epithelialized colonies were rare for C-iPSCs, which bore mesenchymal morphologies (Fig. 1C,D). Some clones reverted to mesenchymal morphologies upon expansion (Fig. 1E). As the emerging epiblasts manifested as epithelialized structures *in vivo* and the epithelialization of mesenchymal cells during reprogramming results from TET activities, we aborted mesenchymal clones for both C-iPSCs and T-iPSCs (Fig. 1B,E). Finally, we established 25 C- iPSC and 145 T-iPSC clones (from 140 and 420 colonies, respectively) (Fig. 1F). A vector-free threshold of 0.03 was employed to exclude clones with episomal vector genome integration (Supplemental Fig. S1C), yielding 46 vector-free clones (7 C-iPSCs and 39 T-iPSCs) from four independent parallel inductions.

**Figure 1.**
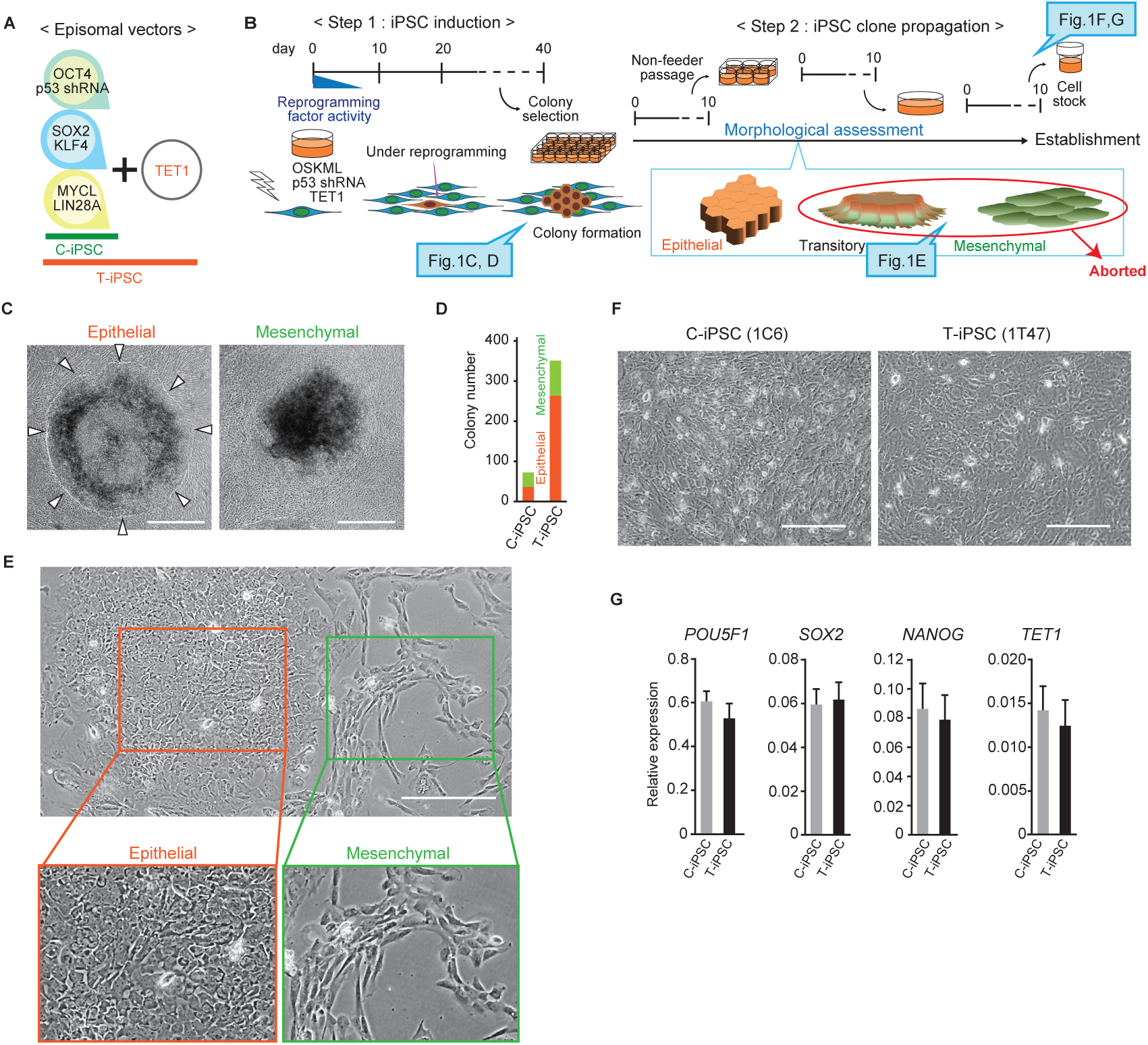
hiPSC-production pipelines. *(A)* Episomal vector combinations for producing C- iPSCs and T-iPSCs. OCT4, an alias for POU5F1. *(B)* Timeline and procedure for hiPSC production. OSKML: OCT4, SOX2, KLF4, MYCL, and LIN28A. *(C)* Characteristic morphologies of emerging hiPSC colonies in Step 1. Arrowheads indicate epithelia-like colony edges. Scale bar = 50 μm. *(D)* Colony numbers with epithelial or mesenchymal morphologies (n = 3). *(E)* A subconfluent culture during the expansion of a colony in Step 2. Scale bar = 100 μm. *(F)* Established C-iPSC and T-iPSC clones. Scale bar = 50 μm. *(G)* Expression of pluripotency markers and *TET1*. Data are presented as the mean ± standard deviation (C-iPSC, n = 7; T-iPSC, n = 39).

To characterize the hiPSC clones at the earliest point, we quantitatively analyzed the endogenous expression of “pluripotency-associated” factors in all clones. We observed no noticeable difference between C-iPSCs and T-iPSCs in expression levels, marker-positive ratios, or sex variance for *NANOG*, *POU5F1,* and *SOX2,* indicating the overall uniformity of the hiPSC clones obtained, and all clones conformed to the qualitative criteria for pluripotency (Fig. 1G; Supplemental Figs. S1D, S2A). By contrast, *KLF4* markedly varied across clones (Supplemental Fig. S2A). Despite the addition of TET1 during reprogramming, T-iPSC clones did not show higher endogenous *TET1* levels than C-iPSC clones (Fig. 1G).

### Interclonal heterogeneity in the differentiation capabilities of hiPSCs

To set a tentative threshold for hiPSC differentiation capability, we chose the default model of neural induction (Fig. 2A). According to this evolutionarily conserved model, dissociated early embryonic cells default to neural induction when incubated in a developmentally neutral culture medium devoid of signaling molecules (Munoz-Sanjuan and Brivanlou 2002; Patani et al. 2009). We selected well-expandable C-iPSCs (1C6) and T-iPSCs (1T35 and 1T47) and pilot- differentiated them. For 1T35 and 1T47, neural differentiation peaked on day 12 of default differentiation, and the expression of PAX6 and SOX1 was maintained until day 18 (Fig. 2B; Supplemental Fig. S2B) (Chambers et al. 2009; Zhang et al. 2010), but 1C6 cells failed to express neural markers and regained pluripotency markers, indicating a defect in pluripotency exit (Supplemental Fig. S2B). We noticed substantial cavities in 1C6-derived spheres, whereas 1T47 produced solid spheres (Fig. 2B). Thus, hiPSCs exhibited considerable differentiation variegation across clones. However, some hiPSC clones can exhibit quasi-total differentiation toward the neural lineage comparable to hESCs (Patani et al. 2009).

**Figure 2.**
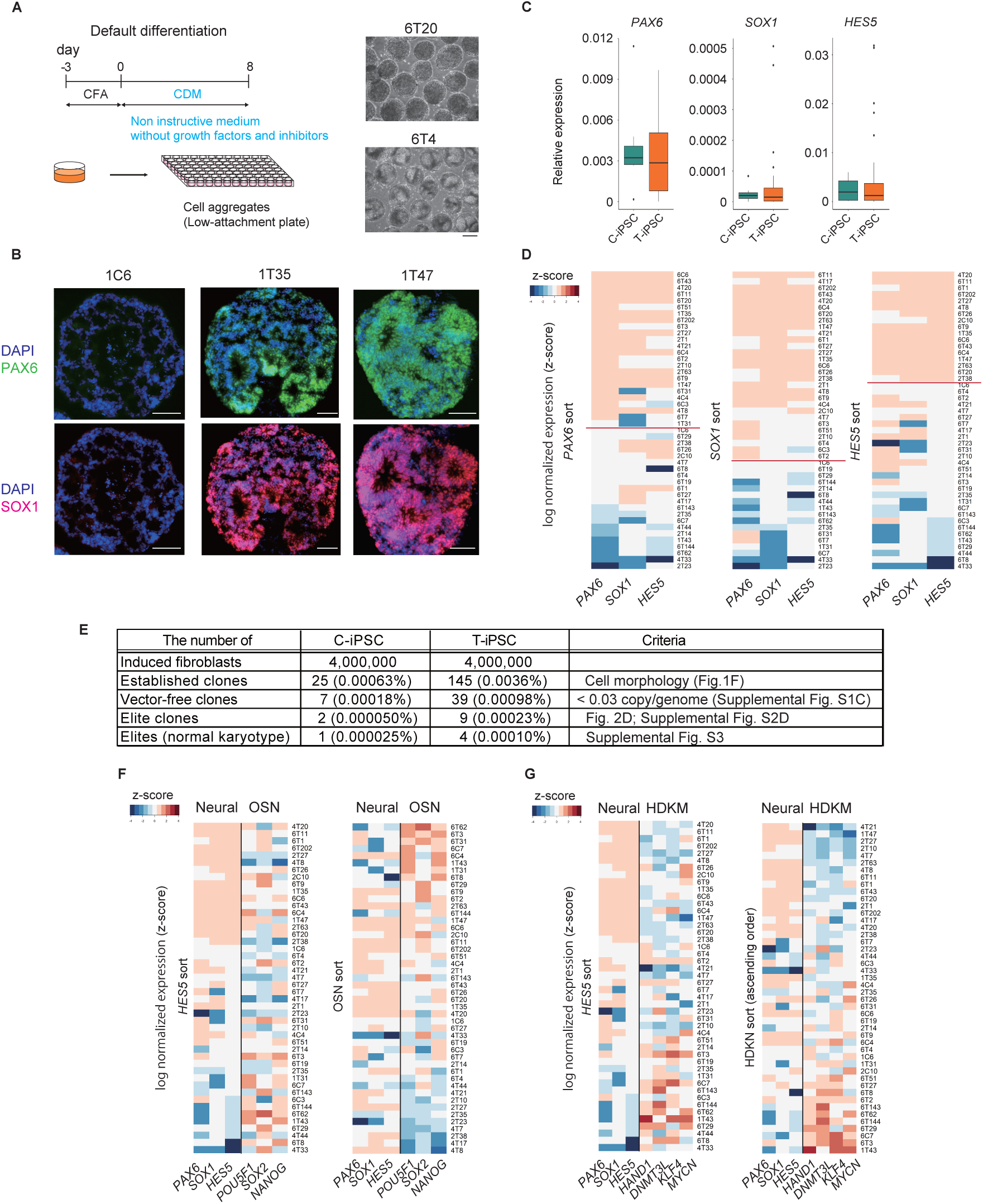
Interclonal differences in differentiation capability between hiPSC clones. *(A)* Schematic representation of the default differentiation timeline and procedure. Clones with high-differentiation capability formed solid spheres of uniform size, whereas clones with low differentiation capability had sac-like structures. Scale bar = 100 μm. *(B)* Representative immunostaining images with DAPI (blue), PAX6 (green), and SOX1 (red) on day 18 of default differentiation. Scale bars = 100 μm. *(C)* Box plots showing the relative expression levels of neural markers on day 8 (C-iPSC, n = 7; T-iPSC, n = 39). *(D)* Heatmap of the relative expression levels of neural markers on day 8. Expression levels were normalized to *GAPDH*, log- transformed, and z-score scaled. Lines indicate tentative thresholds between default-elite and nonelite clones. The rows are sorted by the expression levels of each neural marker. *(E)* Summary of the number of cells (ratio) meeting our tentative criteria in each iPSC production process. *(F)* Heatmap showing relative expression of neural and pluripotency-related markers on day 8. The rows are sorted by the expression levels of *HES5* (left) and the expression levels of pluripotency-related markers (right). O, *OCT4*; S, *SOX2*; and N, *NANOG*. *(G)* Heatmap showing relative expression of neural markers and genes negatively correlated with neural markers on day 8. The rows are sorted by the expression levels of *HES5* (left) and by the expression levels of genes negatively correlated with neural markers (right). H, *HAND1*; D, *DNMT3L*; K, *KLF4*; M, *MYCN*.

To select high-quality hiPSC clones, we investigated each clone using default differentiation. We evaluated expression on day 8 when neural differentiation efficiency diverged (Supplemental Fig. S2B). In addition, we analyzed *HES5* expression (Tchieu et al. 2017). Substantial clones exhibited low differentiation potential, but T-iPSCs contained high-potential outliers (Fig. 2C). *NANOG* (Loh and Lim 2011) and *TBXT* (Sasaki et al. 2016) correlated negatively with neural markers and were used to disqualify clones (Supplemental Fig. S2C,D). We selected clones that outperformed 1C6 in all tests (Fig. 2D) as “default-elite” clones (karyotypes shown in Supplemental Fig. S3).

A retrospective analysis confirmed that the copy number of residual vectors below the set standard of a vector-free threshold (0.03 vector/genome) did not correlate with neural marker expression following default differentiation (Supplemental Fig. S2E). Based on these criteria, only 0.00005% of fibroblasts used for C-iPSC induction acquired the default differentiation capability, a standard that is >100 times more stringent than another report (Fig. 2E) (Okita et al. 2011). TET1-assisted reprogramming increased efficiency to 0.0002%—a fourfold increase in the production of hiPSC clones with enhanced differentiation potential.

### Transcriptome features affecting differentiation efficiency

To link undifferentiated gene expression profiles to differentiation potential, we measured gene expression before (day 0) and after (day 8) default differentiation. We found no gene positively correlated with due differentiation capability (*HES5*/*SOX1*), including classical pluripotency genes (Supplemental Fig. S2C). Conversely, *KLF4* and *MYCN* correlated negatively with neural markers (Supplemental Fig. S2C). Because KLF4 is a factor in differentiation resistance (Kim et al. 2011; Lister et al. 2011; Ohnuki et al. 2014), we investigated KLF4-related factors (DNMT3L and NLRP7) (Cacchiarelli et al. 2015) and HAND1 (Yang et al. 2021). We found that these factors (collectively HDKM genes) negatively correlated with neural markers and *HAND1* positively correlated with aberrant markers (*BMP4* and *NANOG*) (Supplemental Fig. S2C). Clone-by-clone inspection of individual clones confirmed the unrelatedness of classical pluripotency factors to differentiation capability (Fig. 2F). By contrast, clones expressing low *HDKM* in the undifferentiated state tended to express higher levels of the three neural markers (Fig. 2G).

### Neuronal differentiation beyond default neurogenesis

To evaluate differentiation potential beyond the early neural stages assessed by default differentiation, we selected the precursors of the ventral midbrain (VM). We modified a published protocol (Steinbeck et al. 2015) and achieved high efficiency in inducing FOXA2 and LMX1A—VM precursor markers—using the default-elite clones (Andersson et al. 2006). Differentiation efficiency was the highest for 1T47 (Fig. 3A,B; Supplemental Fig. S4A–C). However, some cells lacked LMX1A, NKX6-1, or PAX6, which are markers of VM precursors (Duan et al. 2013). The proportion of these “undefined” cells, presumably induced to cells outside the VM, was significantly higher for 6C6 than for 1T47 (Fig. 3C; Supplemental Table S1). Furthermore, the neuronal dendrites derived from 1T47 were longer than those from 6C6 (Fig. 3D,E).

**Figure 3.**
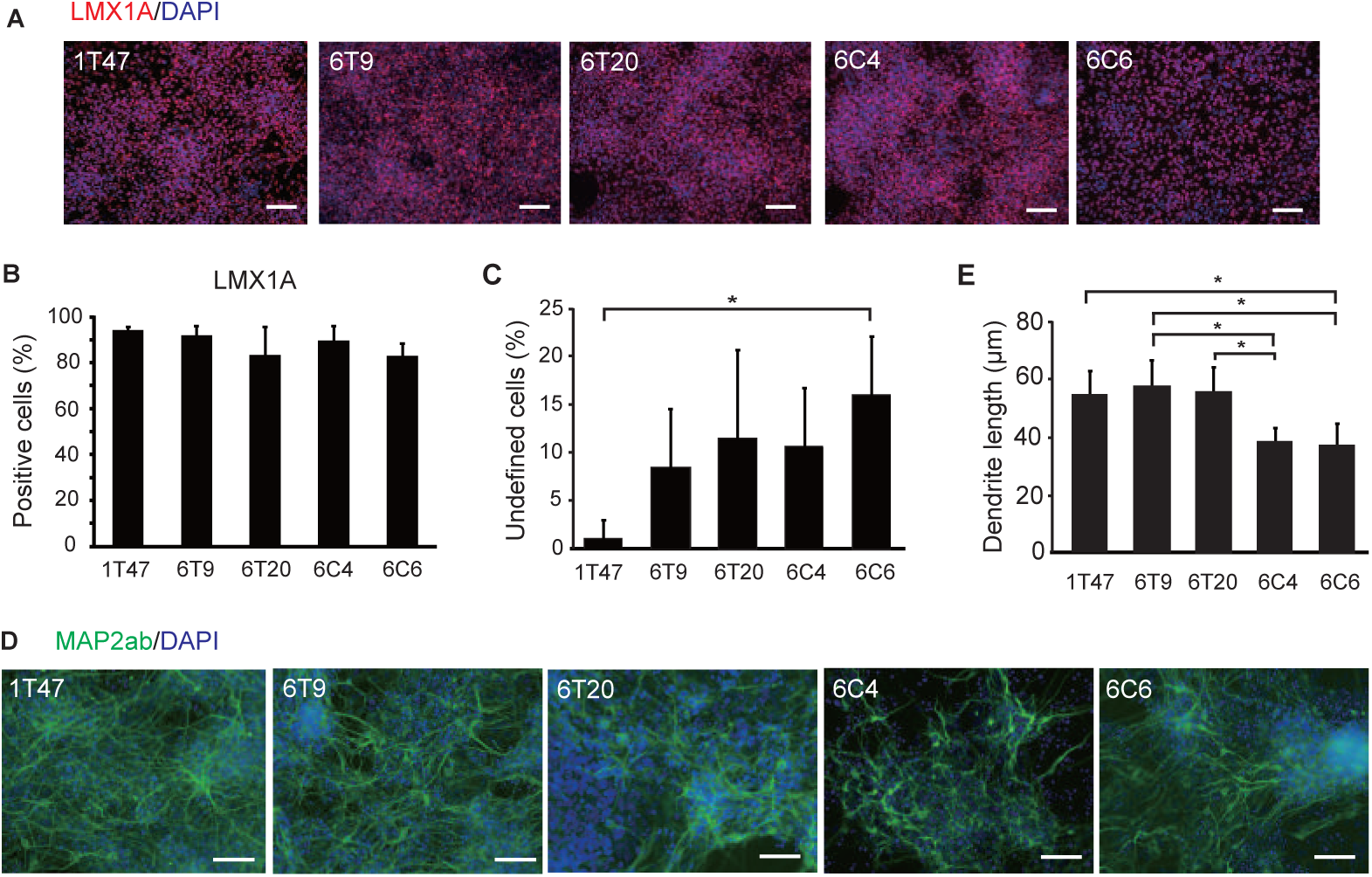
Rationale-based induction of ventral midbrain (VM) progenitors. *(A)* Immunostaining of the hiPSC-derived VM progenitor marker LMX1A (red) with DAPI (blue) on day 19 for default-elite clones. Scale bars = 50 μm. *(B)* Ratio of LMX1A in each clone. *(C)* Ratio of undefined cells (related immunostainings in Supplemental Fig. S4C). *(D)* Immunostaining images with DAPI (blue) and MAP2ab (green) on day 29. Scale bars = 50 μm. *(E)* Lengths of MAP2ab-positive dendrites. Values are presented as mean ± standard deviation (n = 3–4). \**P* < 0.05; One-way analysis of variance and Tukey’s multiple-comparison tests.

### Interclonal heterogeneity of hiPSC clones

We tentatively designated 1T47, with elevated differentiation potentials, as the reference clone. We aimed to identify the distinguishing features unique to 1T47 (T-iPSC^1T47^) compared with other default-elite clones. We selected 6C6 (C-iPSC^6C6^) as our best C-iPSC clones (6C4 was aneuploid; Supplemental Fig. S3). Bulk RNA-sequencing (RNA-seq) analysis identified differentially expressed genes (DEGs) between these clones (Fig. 4A). Gene ontology (GO) analysis indicated few GO terms associated with DEGs expressed at higher levels in T-iPSC^1T47^ (T-iPSC^1T47^-high-DEGs) (Fig. 4B). C-iPSC^6C6^ expressed genes of mesenchymal traits (*TWIST1*, *MYL7, ACTC1, ANXA1*, *VIM*, and *BMP4*) (Fig. 4B,C) (Pham et al. 2022) and genes related to aging, such as cell cycle inhibitors (*CDKN1A/2A/2B*) and senescence-associated secretory phenotype (SASP: *GDF15, IL6, IL1A,* and *IGFBP7*) (Fig. 4C) (Kumari and Jat 2021).

**Figure 4.**
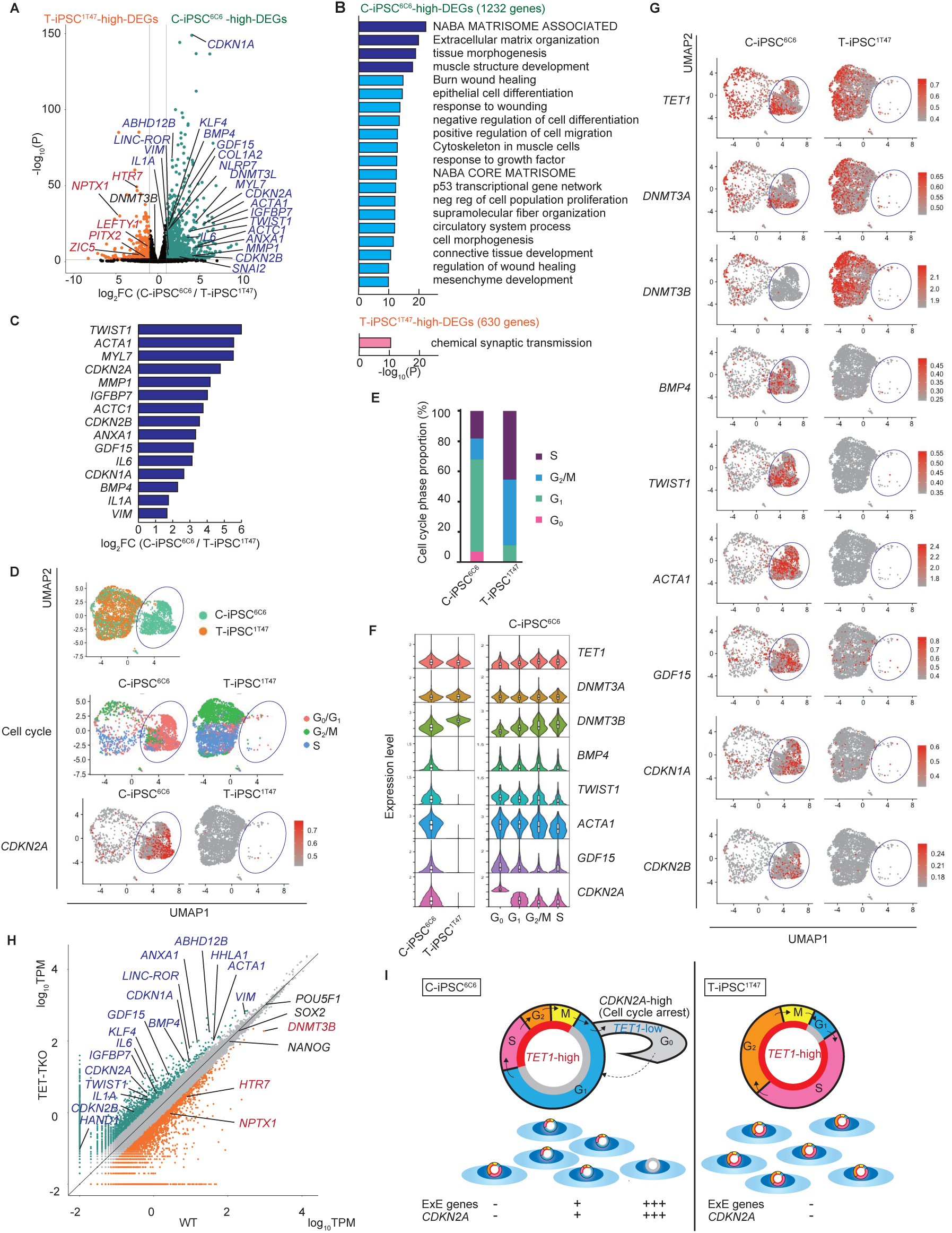
Characterization of interclonal and intraclonal heterogeneities in C-iPSCs. *(A)* DEGs between C-iPSC^6C6^ and T-iPSC^1T47^. Green and orange dots indicate genes with a log2(fold change (FC)) of >1.0 (adjusted *P* value < 0.05). *(B)* GO terms of the DEG sets between C- iPSC^6C6^ and T-iPSC^1T47^. GO terms with −log10(P) >10 are listed. *(C)* Bar plots of relative gene expression of mesenchyme-associated and aging-associated C-iPSC^6C6^-high-DEGs. *(D)* Uniform Manifold Approximation and Projection (UMAP) plot showing C-iPSC^6C6^ and T- iPSC^1T47^ scRNA-seq data, showing the distribution (upper panel) and cell cycle phase distribution (middle panel) as well as *CDKN2A* gene expression (lower panel). Blue circles indicate the C-iPSC^6C6^-specific population. *(E)* Proportions of cell cycle phases in C-iPSC^6C6^ and T-iPSC^1T47^. *(F)* Violin plots showing the expression levels of the regulatory genes for DNA methylation, extraembryonic-associated genes, and aging-associated genes. Gene expression levels in C-iPSC^6C6^ and T-iPSC^1T47^ (left) and at each cell cycle phase in C-iPSC^6C6^ (right). *(G)* Single-cell expression of DNA methylation-, extraembryonic- and aging-associated genes in C-iPSC^6C6^ (left) and T-iPSC^1T47^ (right). Blue circles indicate the C-iPSC^6C6^-specific population. *(H)* Comparison of gene expression between wild-type hESC HUES8 (WT) and TET1/2/3 triple-knockout cells (TET-TKO). Gene expression was plotted as log10(TPM) in WT and TET- TKO. TPM: Transcripts per million. *(I)* The cell cycle affects *TET1* and extraembryonic (ExE) gene expression. C-iPSC^6C6^ exhibits a pronounced cell cycle arrest in the G0/G1 phase, whereas T-iPSC^1T47^ shuttles between S and G2/M. G0/G1 cells express lower levels of *TET1* (*TET1*- low/*CDKN2A*-high) and are the source of ExE gene expression.

Next, we evaluated the versatility of mesenchymal-related and aging-associated gene expression across hiPSC clones. Reanalysis of array data from 49 hiPSC clones revealed significant variation in mesenchymal-related and aging-associated gene expression (Supplemental Fig. S5A,B,D) (Koyanagi-Aoi et al. 2013). Differentiation-defective markers (Ohnuki et al. 2014) were examples of substantial interclonal expression heterogeneities (Supplemental Fig. S5C,D). These results suggest that interclonal heterogeneity is common across hiPSC clones and not unique to our selected reference C-iPSC^6C6^ clone.

### Mesenchymal gene noise in C-iPSCs is related to extraembryonic lineages

The mesenchymal propensity was surprising because hiPSCs are supposedly equivalent to postimplantation epiblasts—an epithelialized tissue (Nakamura et al. 2016). Because obtaining human embryonic gene profiles from implantation to gastrulation is difficult, we used single-cell RNA-seq (scRNA-seq) data of peri-gastruloids (Liu et al. 2023) to determine lineages that express mesenchymal genes during human development. Peri-gastruloids encompass developmental trajectories from the epiblast to the definitive mesoderm/endoderm and extraembryonic lineages (Supplemental Fig. S6A–D). We integrated peri-gastruloid scRNA-seq data with data obtained from T-iPSC^1T47^, C-iPSC^6C6^, and a standard hESC H9 (Messmer et al. 2019). Although the mesenchymal markers (*ACTC1*, *BMP4*, *HAND1*, *MYL7*, *TWIST1*, and *VIM*) are widely recognized as mesodermal markers, cells expressing these genes in the peri-gastruloids did not coexpress the definitive mesodermal markers, *MESP1/2* and *MIXL1* (Supplemental Fig. S6C,D) (Liu et al. 2023). Notably, this population exclusively expressed *ANXA1*, a human extraembryonic mesoderm marker (Supplemental Fig. S6D) (Pham et al. 2022). T-iPSC^1T47^ and hESC H9 cosegregated close to the epiblast clusters, whereas C-iPSC^6C6^ additionally scattered within the extraembryonic mesoderm (Supplemental Fig. S6A,E). These data indicate that a subpopulation of C-iPSC^6C6^ is akin to the extraembryonic mesoderm rather than the definitive mesoderm (Fig. 4C; Supplemental Fig. S6C,D). To consolidate the developmental contexts of hPSCs, we referred to cultured monkey blastocysts up to the gastrulation stage (Supplemental Fig. S7A) (Yang et al. 2021). Consistently, C-iPSC^6C6^-high-DEGs were expressed in the amnion or extraembryonic mesoderm and low in the definitive mesodermal lineages (Fig. 4C; Supplemental Fig. S7B). These results indicate that C-iPSC^6C6^ expresses extraembryonic mesodermal/mesenchymal rather than embryonic mesodermal genes.

### Intraclonal heterogeneities in C-iPSCs

scRNA-seq analyses revealed a proportion of C-iPSC^6C6^ cells with halted cell cycles (Fig. 4D). We defined cells with higher *CDKN2A* expression and lower *MKI67* expression as being in the G0 phase (Supplemental Fig. S8A). Based on this, 7% of C-iPSC^6C6^ and no T-iPSC^1T47^ were in the G0 phase (Fig. 4E). *TET1*, *DNMT3B*, and *SOX2* expression was lower in the G0 phase (Fig. 4D,F,G; Supplemental Fig. S8B–D). Notably, C-iPSC^6C6^ cells in the G0 phase highly expressed extraembryonic genes (Fig. 4D,F,G; Supplemental Fig. S8B–D). Given the lower expression of *TET1* in cells expressing cell cycle inhibitors and extraembryonic markers, we reanalyzed the RNA-seq data for *TET1/2/3*-triple-knockout hESCs (TET-TKO) (Charlton et al. 2020). TET-TKO exhibited increased expression of cell cycle inhibitors, extraembryonic markers, and differentiation-defective markers, as in C-iPSC^6C6^ (Fig. 4H; Supplemental Fig. S8E). Collectively, a subpopulation of C-iPSCs in the G0 phase had low *TET1* expression, exhibiting mesenchymal/extraembryonic gene noise (Fig. 4I).

We evaluated whether the observed differentiation superiority of T-iPSC^1T47^ (Fig. 3) is a trade-off for their inability to differentiate into extraembryonic lineages. We proactively differentiated T-iPSC^1T47^ by adding BMP4, which induces extraembryonic cells (Chhabra et al. 2019; Pham et al. 2022). *BMP4*, a marker of extraembryonic cells, was self-induced at a higher level in T-iPSC^1T47^ than in C-iPSC^6C6^ (Supplemental Fig. S9A). BMP4 induced *TWIST1* and *ANXA1* expression in T-iPSC^1T47^, indicating its potential to differentiate into extraembryonic lineages (Supplemental Fig. S9A). T-iPSC^1T47^ exhibited higher differentiation efficiency into somatic mesendodermal lineages when C-iPSC^6C6^ retained undifferentiated pluripotency markers (Supplemental Fig. S9B,C). These results indicate that T-iPSCs are not biased toward the neural lineage and can differentiate into the three germ layers and extraembryonic tissues— validating human pluripotency.

### TET1 insufficiency leads to hypermethylation of poised extraembryonic gene enhancers

To determine the effect of TET1 insufficiency, we compared DNA methylomes of wild-type (WT) hESCs, TET-TKO (Charlton et al. 2020), fibroblasts, T-iPSC^1T47^, and C-iPSC^6C6^ using whole genome bisulfite sequencing (WGBS). We examined differentially methylated regions (DMRs) and identified 4769 C-iPSC^6C6^- and 1244 T-iPSC^1T47^-hypermethylated DMRs (hyper-DMRs). DMRs were observed around the C-iPSC^6C6^-high-DEG loci of extraembryonic markers, aging-related markers, and *KLF4* (Fig. 5A; Supplemental Fig. S10A). Although C- iPSC^6C6^-hyper-DMRs were enriched in both promoters and gene bodies, T-iPSC^1T47^-hyper- DMRs were enriched in promoters (Fig. 5B). C-iPSC^6C6^-hyper-DMRs were associated with genes involved in developmental morphogenesis and epithelial-mesenchymal plasticity (Supplemental Fig. S10B) (Haerinck et al. 2023). Although DNA hypermethylation silenced gene expression in T-iPSC^1T47^, it facilitated gene expression in C-iPSC^6C6^ (Fig. 5C; Supplemental Table S2) (Jones 1999). This paradox was resolved by analyzing the Polycomb repression mark, H3K27me3. Both C-iPSC^6C6^- and T-iPSC^1T47^-hyper-DMRs attracted H3K4me1, H3K4me3, H3K27ac, and H3K27me3 in hPSCs (Supplemental Fig. S10C,D). Especially, C-iPSC^6C6^-hyper-DMRs exhibited significant enrichment of H3K4me1 and H3K27me3 in hESCs (Fig. 5D). The regions co-occupied with a putative enhancer mark (H3K4me1) and the Polycomb repression mark (H3K27me3) are categorized as “poised enhancers” (Rada-Iglesias et al. 2011). We found gene body-embedded poised enhancers intersecting C-iPSC^6C6^-hyper-DMRs in hPSCs (Fig. 5A; Supplemental Fig. S10A). In a reference hESC clone, H3K27me3 was significantly enriched in the C-iPSC^6C6^-hyper-DMRs associated with the C-iPSC^6C6^-high-DEGs (Fig. 5E; Supplemental Fig. S10E). Notably, C- iPSC^6C6^-hyper-DMRs associated with C-iPSC^6C6^-high-DEGs were significantly enriched in putative enhancer regions (Fig. 5F; Supplemental Fig. S10F). Hypermethylation of poised enhancers could cause target gene derepression because DNA methylation repels Polycomb repressive complex 2 (PRC2) (Blackledge et al. 2015). These results suggest that TET1 insufficiency leads to the hypermethylation of poised enhancers and the paradoxical derepression of the related genes. Of the 1232 C-iPSC^6C6^-high-DEGs, 350 were upregulated in PRC2 component EZH2-knockout hESCs (EZH2-KO) (Fig. 5G,H) (Collinson et al. 2016). We identified 127 genes commonly overexpressed in C-iPSC^6C6^, EZH2-KO, and TET-TKO (Fig. 5H; Supplemental Table S3). GO analysis of these common genes showed enrichment of epithelial-mesenchymal transition-related terms (Fig. 5I). These findings suggest that the hypermethylation of poised enhancers in C-iPSC^6C6^ resulting from TET1 insufficiency may be the cause of the derepression of cell cycle inhibitors and extraembryonic mesenchymal genes by evading Polycomb-mediated repression and the ensuing morphological changes.

**Figure 5.**
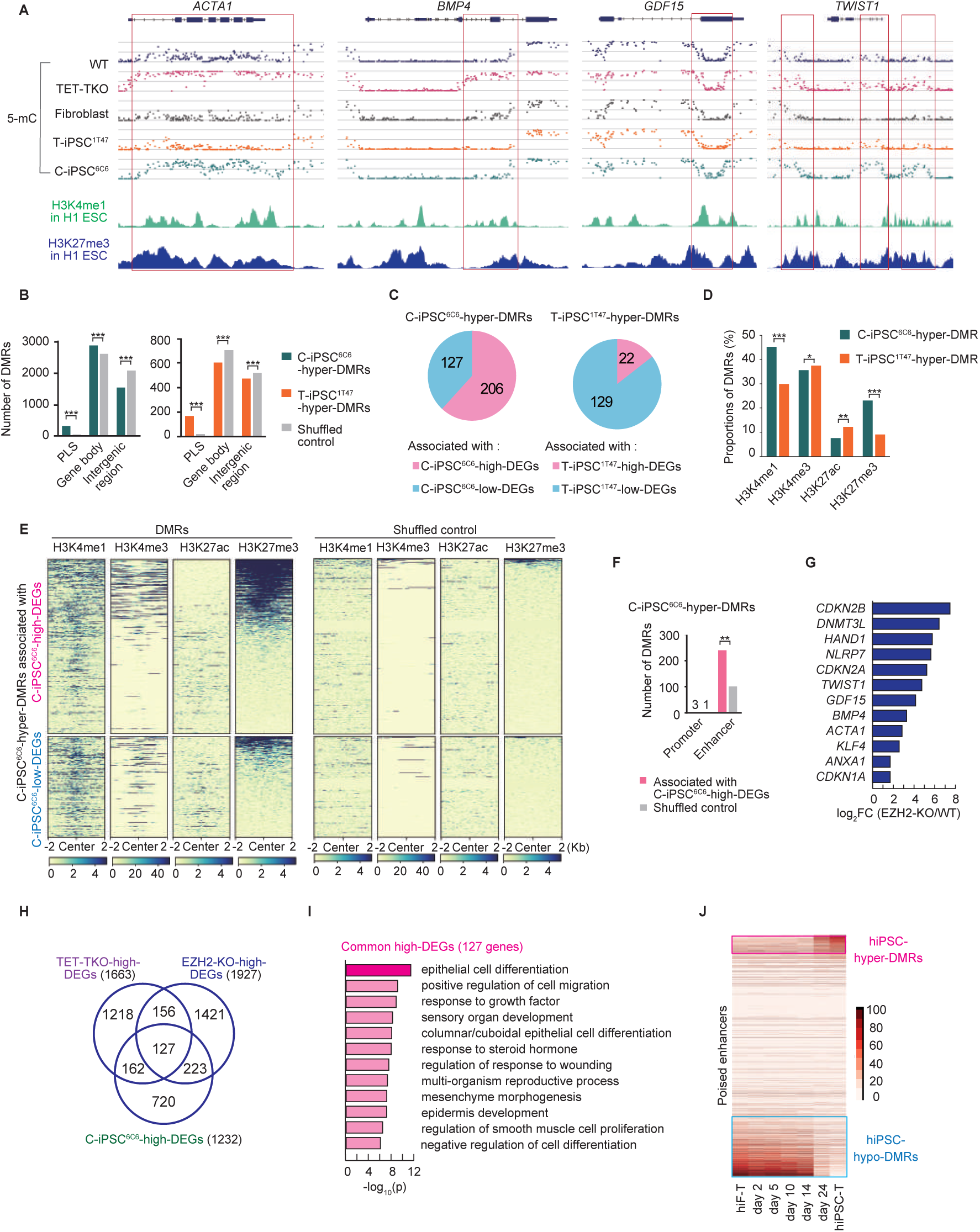
Hypermethylation of poised enhancers on extraembryonic gene loci. *(A)* WGBS (5- mC) and ChIP-seq data of extraembryonic genes. DMRs between C-iPSC^6C6^ and T-iPSC^1T47^ are indicated by red rectangles. WT, hESC HUES8; TET-TKO, TET1/2/3-triple-knockout HUES8. *(B)* Bar plots showing the distribution of hyper-DMRs in the genomes of C-iPSC^6C6^ and T-iPSC^1T47^. PLS; promoter-like signature. *(C)* The proportion of high- and low-DEGs associated with DMRs between C-iPSC^6C6^ and T-iPSC^1T47^. (*P* < 1e-100; chi square test). *(D)* Proportions of hyper-DMRs overlapping with histone modifications in hESC H1. *(E)* Heatmap of ChIP-seq signals of histone modifications (hESC H1) in C-iPSC^6C6^-hyper-DMRs associated with the DEGs between C-iPSC^6C6^ and T-iPSC^1T47^. The rows are sorted in the order of H3K27me3-densities. *(F)* Bar plots showing the number of C-iPSC^6C6^-hyper-DMRs associated with C-iPSC^6C6^-high-DEGs on *cis*-regulatory elements. *(G)* Bar plots of selected common gene expression levels to C-iPSC^6C6^-high-DEGs and EZH2-KO-high-DEGs. WT, hESC H9; EZH2- KO, EZH2-knockout H9. *(H)* Common high-DEGs across C-iPSC^6C6^, TET-TKO, and EZH2- KO. *(I)* GO terms of common high-DEGs across C-iPSC^6C6^, TET-TKO, and EZH2-KO. *(J)* Heatmap of the DNA methylation profile at poised enhancers during human reprogramming (Cacchiarelli et al. 2015). In the comparison between hiF-T and hiPSC-T, hiPSC-hyper-DMRs and hiPSC-hypo-DMRs are indicated in pink and blue, respectively. hiF-T, TERT- immortalized fibroblast; hiPSC-T, reprogrammed hiPSC. \*\*\**P* < 1e-100, \*\**P* < 1e-20, and \**P* <0.01; chi square test.

### Aberrant methylation is acquired during iPSC reprogramming

Most C-iPSC^6C6^-hyper-DMRs within gene bodies were hypomethylated in fibroblasts (Fig. 5A), suggesting that these regions acquired methylation during reprogramming (Ma et al. 2014; Roost et al. 2017; Buckberry et al. 2023). We reanalyzed methylome data from conventionally reprogrammed iPSCs (Cacchiarelli et al. 2015) and found 1732 autosomal enhancers that were poised in hPSCs and acquired *de novo* DNA methylation during reprogramming (Fig. 5J). Approximately 70% of the enhancers were within gene bodies, and *de novo* methylated poised enhancers were similarly distributed (Supplemental Fig. S10G). These results indicated that many poised enhancers in hiPSCs acquired *de novo* methylation in the gene body as the sole consequence of reprogramming. We assessed their expression during reprogramming because gene-body methylation primarily correlates with gene expression in the presence of DNMT3B (Jeziorska et al. 2017). As expected, these extraembryonic genes were transiently expressed during reprogramming (Supplemental Fig. S10H) (Liu et al. 2020). The DNA methylation pattern in hPSCs is constantly regulated by counteracting methylating (DNMT3A/B) and demethylating (TET1) activities (Charlton et al. 2020). This balance may be disrupted in a subset of cells with insufficient TET1 and DNMT3B activity during reprogramming (Supplemental Fig. S1A,B) and in the G0 phase in hiPSCs (Fig. 4F).

## Discussion

Our default differentiation survey showed that the expression of individual pluripotency-associated markers not only did not positively correlate to differentiation results but also negatively correlated to neural induction (*KLF4*) in some instances. The “precarious balance model” argues that pluripotency factors act as lineage specifiers, and the balance of these factors is important for maintaining the undifferentiated pluripotent state (Loh and Lim 2011). Therefore, the individual expression of pluripotency markers should be interpreted cautiously and not be used alone to validate the pluripotency of the relevant PSC clone. Notably, no markers showing a positive correlation with differentiation capabilities were identified. This observation supports the idea that variations in the quality of pluripotent stem cells are reflected in the aberrant upregulation of lineage-specific, typically silenced genes. Such derepression is likely driven similarly to the epigenetic drift proposed in the Information Theory of Aging (Lu et al. 2023).

Default differentiation is not merely a single lineage differentiation test for neural induction. Human default differentiation may discern the propensity between somatic/embryonic versus extraembryonic differentiation. In addition to its allowance for quantitative measurements, the default test has practical advantages. Compared to *in vivo* teratoma formation, it is a short-term test, making it less labor-intensive, and the defaultness helps to provide robustness and reproducibility, making it suitable for a wide range of laboratory settings.

The expression of extraembryonic markers in hPSCs evokes a different scenario to the primed pluripotency proposed for mouse PSCs, where cells are halted before differentiation of the germ lineage (Smith 2017). Considering the wide occurrence of *CDKN2A* and extraembryonic gene expression across hiPSCs, a subset of cells residing in the G0 phase might be common. We found that cells in the G0/G1 phase had lower *TET1* expression (Fig. 4F), which may have led to the hypermethylation of the poised enhancers (Fig. 5D–F). In turn, the hypermethylation of nearby poised enhancers may have caused the derepression of extraembryonic genes (Blackledge et al. 2015). Consistently, in extraembryonic tissue, hyper-DMRs frequently overlapped with PRC2-regulated genes in mouse pluripotent cells (Smith 2017) and EZH2-KO exhibited extraembryonic gene dysregulation (Collinson et al. 2016). *BMP4* expression is commonly found across hiPSCs (Buckberry et al. 2023) and may assist in an autocrine/paracrine manner to aggravate aberrant extraembryonic gene expression. This highlights the importance of standardization when optimizing target cell derivation in regenerative medicine because the outcome would be affected by the selection of appropriate hiPSC clones and the combined differentiation protocol, which should be established using a standardized PSC clone. This may explain the current scarcity of universally applicable differentiation protocols.

The bystander effect of secreted proteins, such as BMP4, from cell cycle-arrested cells is strongly reminiscent of cellular senescence. Indeed, G0 cells exhibited a senescence-associated secretory phenotype (SASP) (Fig. 4C). The appearance of G0/senescent cells was unexpected in stem cells. Considering that aging-related hypermethylated gene loci are Polycomb domains (Lu et al. 2023; Moqri et al. 2024), a wide survey of the “ages” of various hPSCs would be informative.

This also raises the possibility that these epigenetic controls are reciprocally linked to cell cycle regulation. TET1 and PRC2 establish bivalent chromatin, a cardinal feature of pluripotency (Verma et al. 2018). Cells that fail to achieve adequate pluripotency or bivalency during development may be “outcompeted” through CDKN2A expression, leading to premature senescence and preventing their contribution to further development. This developmental safeguard mechanism may cause suboptimal hPSCs to exhibit heterogeneity.

This study has some limitations. We could not decipher how the ectopic TET1—during reprogramming—directed homogeneous endogenous *TET1* expression or mechanistically conferred differentiation advantages to T-iPSCs partly as complete reprogramming is a rare event (one T-iPSC^1T47^ clone/11 default-elite clones/46 integration-free clones/398 epithelialized colonies). As the episomal vectors used in this study exhibit only transient expression (approximately 7 days) and were integrated in half of our established clones, introducing TET1 as synthetic RNA multiple times during reprogramming may enhance the efficiency of hiPSC clones with attenuated heterogeneities. Currently, we are working on obtaining female T-iPSCs, which may exhibit equivalent capabilities to the male 1T47 clone, as this would allow further investigations related to female cell-specific events (oogenesis, X-chromosome inactivation). In this study, we proposed the derepression of PRC2 target genes as a mechanism underlying the inconsistent differentiation potential of iPSCs. While our sample size is limited, we hope our findings motivate a consortium effort to validate whether the differentiation potential of iPSCs globally correlates with the DNA methylome of poised enhancers.

In summary, we have developed a new hiPSC induction system coupled with default differentiation screening, showing the quasi-total induction of VM progenitors from T-iPSCs. Our results strongly suggest that the currently proposed method for creating hiPSCs induces heterogeneities due to TET1 insufficiency during and after reprogramming. Developing strategies to avoid these heterogeneities will be critical for advancing the practical application of hiPSCs, and T-iPSCs may be a promising avenue for accomplishing this.

## Acknowledgments

We thank Y. Moriyama, Y. Tokuzawa, S. Funayama, R. Hamada, and H. Kawakami for earlier contributions and technical assistance. This study was supported by Zeon Corporation, SATAKE MultiMix Corporation, Yokogawa Electric Corporation, SyntheticGestalt, and AMED under Grant Number JP20bk0104090. We thank Perfect C. (Edanz) and Laura J (Enago) for editing.

## Author contributions

H. Kato conceived the project. M. M., K. H-K., and H. Kato wrote the original draft. M. M., K. H-K., M. K., and H. Kato performed the culture and differentiation experiments and analyzed the data. K. H-K. and M. K. analyzed the sequencing data and performed the bioinformatic analyses under the supervision of H. Kato. M. M., K. H-K., M. K., H. Kiyosawa, Y. K., and H. Kato reviewed, edited, and approved the final version of the manuscript.

## Materials and methods

### hiPSC induction and culture

Human fibroblasts (from male neonatal foreskin, 106-05N; female fetal fibroblasts, 106-05F) were purchased from Cell Applications Inc., and cultured in Dulbecco’s modified Eagle’s medium (DMEM; Nacalai Tesque) supplemented with 10% fetal bovine serum (FBS; Hyclone), 100 μg/mL L-ascorbic acid 2-phosphate trisodium salt (L-AA; FUJIFILM Wako), and 1× penicillin–streptomycin (P/S; Gibco). hiPSCs were induced using episomal vectors (Supplemental Table S4) (Okita et al. 2011). To establish T-iPSCs, a TET1 episomal vector (comprising complementary DNA encoding the human *TET1* open-reading frame, cloned into the pCXLE vector) was used with the three episomal vectors upon reprogramming. One µg of each vector was transfected into approximately 1 × 10^6^ low passage fibroblasts using Human Dermal Fibroblast Nucleofection Kit and Nucleofector Transfection 2b Device (Amaxa Biosystems [Lonza]), according to the manufacturer’s instructions, and the cells were plated in a human fibroblast medium. The next day, the culture medium was replaced with Essential 8 Medium (E8M; Gibco) supplemented with 100 nM hydrocortisone (Sigma-Aldrich) and 1 × P/S. E8M, which contains TGF-β, was selected because of its potentiating effect of activating SMAD2/3 during reprogramming (Ruetz et al. 2017). The medium was changed every other day. E8M without hydrocortisone was used from day 10 of iPSC induction. iPSC colonies were picked up, triturated into E8M supplemented with 10 mM Y-27632 (FUJIFILM Wako), and plated onto STO feeder cells (RIKEN BRC, RCB0536; RRID: CVCL_3420) treated with mitomycin C (FUJIFILM Wako). The culture medium was replaced the following day with KnockOut DMEM/F12 (Thermo Fisher Scientific) supplemented with 20% KnockOut Serum Replacement (KSR), 2 mM GlutaMAX^TM^-I (Glu), 1 × MEM Non-Essential Amino Acids Solution (NEAA) (Gibco), 0.1 mM 2-mercaptoethanol (2-ME), 5 mM NaOH (FUJIFILM Wako), 5 ng/mL fibroblast growth factor 2 (FGF2; Peprotech), and 1 × P/S. After the colonies had been picked, the medium was changed daily. Grown iPSC colonies were dissociated with 0.01% trypsin/0.1 mM EDTA buffer (trypsin/EDTA; Nacalai Tesque) and plated onto 5 μg/mL vitronectin (Invitrogen)-coated 6-well plates in KnockOut DMEM/F12 supplemented with 20% KSR, 2 mM Glu, 1 × NEAA, 0.1 mM 2-ME, 15 ng/mL FGF2, 10 ng/mL Activin A (R&D Systems), 5 mM NaOH, and 1 × P/S (KFA medium) supplemented with 10 mM Y-27632. The culture medium was replaced daily with KFA medium without Y-27632. KFA medium was routinely used throughout this study to expand and maintain hiPSCs. Clones that reached >90% confluency in a 100-mm dish (Becton Dickinson) without a mesenchymal morphology were considered “established.” One-half of the cells were cryopreserved in a cell stock solution (10% dimethyl sulfoxide [Nacalai Tesque] and 90% FBS). Genomic DNA (gDNA) was obtained from the remaining cells to analyze the genomic integration of the plasmid vectors.

### Setting a threshold for the genomic integration of episomal vectors

gDNA was extracted using DNeasy Blood & Tissue Kit (QIAGEN). Conventional PCR was performed using Ex Taq DNA polymerase (TaKaRa). Quantitative PCR (qPCR) was performed using Thunderbird SYBR qPCR Mix (Toyobo) and CFX Connect Real-Time System (Bio-Rad). The cycle threshold (Ct) values of the episomal vector were normalized against gDNA measured using the CFTR primer set (Supplemental Table S4). The pEP4-SF1 and pEP4-SR1 primers were used to detect episomal vector backbones for conventional PCR (International Stem Cell et al. 2007). For qPCR, the pEP4-SR1 primer was replaced with the pEP4-SR3 primer, which was designed in-house. The template for conventional PCR and qPCR was 50 and 20 ng gDNA, respectively. gDNA extracted from male human dermal fibroblasts was used as a negative control.

### Differentiation of hiPSCs by default differentiation

Pilot experiments using hiPSCs and attempts to derive neural cells from monkey ESCs (Ono et al. 2014) indicated that the KSR-containing culture media are incompatible with default differentiation. Therefore, we gradually changed KSR medium to a chemically defined medium (CDM) before differentiation to facilitate the efficient derivation of neural cells (Ono et al. 2014). KFA-cultured hiPSCs were dissociated with trypsin/EDTA, and 1 × 10^6^ cells were replated onto a vitronectin-coated 100-mm dish in CDM supplemented with 10 ng/mL FGF2, 2 ng/mL Activin A, 3% KSR, and 10 μM Y-27632 (CFA). CDM is composed of Iscove’s modified Dulbecco’s medium/F-12 (Thermo Fisher Scientific), 2 mM Glu, 7 μg/mL Insulin (FUJIFILM Wako), 15 μg/mL transferrin, 450 μM monothioglycerol, 5 mg/mL bovine serum albumin fraction V, 80 nM sodium selenite (Sigma-Aldrich), 200 μM L-AA, 1 × lipid concentrate (Gibco), 1 × NEAA, and 1 × P/S. One-half of the media was carefully replaced with CFA without Y-27632 daily. On day 3, the cells were dissociated with trypsin/EDTA and plated at 500 cells per well in a 96-well low-binding Nunclon Sphera Plate (Thermo Fisher Scientific) in 150 μL of CDM supplemented with 10 mM Y-27632. The day on which the cells were seeded was designated differentiation day 0. Two days after seeding, 100 μL of the medium was removed to avoid aspiration of the spheres, and 150 μL of fresh CDM without Y- 27632 was added. On differentiation day 5, 100 μL of the medium was removed, and 100 μL of fresh CDM was added. The medium was changed every 3 days using the same procedure. The induced aggregates were collected and analyzed on differentiation days 8 and 18.

### Differentiation of ventral midbrain neural progenitor cells

A step-by-step protocol can be found in Supplemental materials and methods. We optimized the timing of adding the reagents mainly involved in WNT signaling because WNT activity initially inhibits neurogenesis but is imperative for coordinating the anterior–posterior axis of the midbrain. Briefly, 100 nM LDN193189 was added from days 0 to 11, 10 μM SB431542 from days 0 to 4, and 10 nM WNT-C59 (C59) from days 0 to 2. Ventralization was initiated by adding 100 ng/mL recombinant sonic hedgehog (C24II) (R&D Systems) and 2 μM purmorphamine (FUJIFILM Wako) from days 3 to 8. Furthermore, 3 μM CHIR99021 (Sigma-Aldrich) was added from days 3 to 12. From differentiation day 13, neural maturation was initiated by adding 10 ng/mL brain-derived neurotrophic factor, 10 ng/mL glial-derived neurotrophic factor (FUJIFILM Wako), 1 ng/mL TGF-β3 (R&D Systems), 200 μM L-AA, 200 μM *N*6,2-*O*-dibutyryladenosine 3, 5-cyclic monophosphate (Sigma-Aldrich), and 5 μM DAPT (Tocris Bioscience). The induced cells were dissociated with Accumax (Innovative Cell Technologies, Inc.) on day 16. Then, 4 × 10^5^ cells/cm^2^ were seeded onto culture slides (BD Biosciences) coated with 15 μg/mL poly-L-ornithine, 2 μg/mL fibronectin (Sigma-Aldrich), and 0.5 μg/cm^2^ laminin. The medium was changed every day from days 0 to 15, and one-half was changed every 2–3 days from day 16 onward.

### Immunohistochemical analysis

Default differentiated spheres were washed with phosphate-buffered saline (PBS; TaKaRa), fixed in 4% paraformaldehyde (PFA; FUJIFILM Wako), embedded in OCT compound, cryosectioned at 10 μm using a cryostat (Leica Biosystems), and collected onto glass slides. Undifferentiated hiPSCs and ventral midbrain precursors were cultured on glass chamber slides. The slides were fixed in 4% PFA and blocked in 5% (v/v) normal donkey serum (Millipore)/0.3% (v/v) Triton X-100 (Nacalai Tesque)/PBS and then incubated overnight with primary antibodies (Supplemental Table S5). The slides were washed and incubated with secondary antibodies and DAPI (Invitrogen) (Supplemental Table S5). Finally, the slides were washed with 0.05% Tween-20/PBS and mounted with an anti-photobleaching mounting medium. The stained cells were imaged using a Biorevo BZ-9000 fluorescence microscope (Keyence).

### Reverse transcription-qPCR analysis

Total RNA was purified from the samples using RNeasy Plus Mini Kit (QIAGEN). More than 50 ng of total RNA was reverse transcribed using ReverTra Ace and oligo-dT primers (Toyobo). qPCR analysis was performed using Thunderbird SYBR qPCR Mix and CFX Connect Real-Time System. Data were assessed using the delta-Ct method and normalized to *GAPDH* expression. Error bars represent the standard deviation of biological replicates. Primer sequences used for qPCR are listed in Supplemental Table S4. Statistical significance was determined using Student’s unpaired *t*-test or one-way analysis of variance and Tukey’s multiple-comparison tests using R software. *P* < 0.05 indicated statistically significant differences.

### Evaluating standards for default-elite clones

Considering the conducted correlation analyses of differentiation markers and differentiation efficiencies between 1C6, 1T35, and 1T47, we set our default-elite thresholds to *HES5*/*GAPDH* >0.002 and *TBXT*/*GAPDH* <0.0001 detected by qPCR.

### RNA-seq processing

RNA-seq data analyses were performed for T-iPSC^1T47^ and C-iPSC^6C6^ cultured under an undifferentiated culture condition (KFA medium). Total RNA was purified using RNeasy Plus Mini Kit, according to the manufacturer’s instructions. Then, library preparation and sequencing were performed by Macrogen Japan Corp. Briefly, RNA-seq libraries were prepared using TruSeq Stranded Total RNA LT Sample Prep Kit with Ribo-Zero Human/Mouse/Rat (Illumina). By harvesting cells at different passage numbers, we generated biological triplicate data for T-iPSC^1T47^ and C-iPSC^6C6^. RNA-seq libraries were sequenced on an Illumina NovaSeq 6000, generating 100-base paired-end reads.

### RNA-seq data analysis

The software and resource data are listed in Supplemental Table S6. RNA-seq reads were aligned to the human reference genome (GRCh38/hg38) using STAR software with default parameters. We calculated transcripts per million mapped reads using RSEM with the default parameters and considered their expression levels. The DEGs between T-iPSC^1T47^ and C- iPSC^6C6^ and between EZH2-KO and WT hESCs were calculated using the R package DESeq2 with the cutoffs of absolute log2(FC) >1 and adjusted *P* value of <0.05. The DEGs between TET-TKO and WT hESCs were calculated using a cutoff of absolute log2(FC) >1. We only considered autosomal chromosomes for analyzing RNA-seq data.

### Isolation and RNA-seq of single cells

scRNA-seq was performed for T-iPSC^1T47^ and C-iPSC^6C6^ cultured in an undifferentiated culture condition (KFA medium). Azenta Life Sciences performed library preparation and sequencing. Briefly, scRNA-seq was performed using the 10 × Chromium platform and Chromium Single- Cell 3′ Reagent Kits (v3 Chemistry), according to the manufacturer’s protocol (10 × Genomics). Libraries were sequenced on DNBSEQ-G400 (MGI) with paired-end sequencing.

### scRNA-seq data analysis

Alignment of reads to the human reference genome (GRCh38/hg38) and preparation of the raw gene expression matrix were conducted using the Cell Ranger pipeline. scRNA-seq integration analysis was performed using the Seurat package. C-iPSC^6C6^ cells and T-iPSC^1T47^ cells were excluded if they contained a portion of mitochondrial reads >20% or had <200 genes detected. The resulting matrix contained 2119 C-iPSC^6C6^ and 3249 T-iPSC^1T47^. Unique molecular identifier counts were normalized, followed by variable gene identification, anchor identification, data integration, and data scaling with the default settings of the Seurat package. Cell cycle phases (G0/G1, S, and G2/M) were determined using Seurat’s default settings. Cells in the G0 phase were selected from the G0/G1 phase subset based on *CDKN2A* expression (>1). For Uniform Manifold Approximation and Projection (UMAP), we used the first 15 principal component analysis dimensions with the default settings using the RunUMAP function. Gene expression profiles were visualized on a UMAP plot using the FeaturePlot function (min.cutoff = q50, max.cutoff = q80). For clustering, we used a resolution of 0.5 with default settings.

To reanalyze scRNA-seq data from peri-gastruloids, we integrated scRNA-seq data from T-iPSC^1T47^, C-iPSC^6C6^, and hESC H9. Cells were excluded if they had <200 genes detected. We randomly selected 200 cells each from the T-iPSC^1T47^, C-iPSC^6C6^, and primed hESC H9 datasets. The resulting matrix contained 200 cells (C-iPSC^6C6^), 200 cells (T-iPSC^1T47^), 200 cells (hESC H9), and 12,008 cells (peri-gastruloids). Unique molecular identifier counts were normalized, followed by variable gene identification with default settings of the Seurat package. For integration with peri-gastruloids, merged hiPSCs, and hESCs in the primed culture condition, we identified anchors using the default settings, except for anchor.features = 1500 and k.filter = 10. Integration and data scaling were performed with the default settings. For *t*-distributed stochastic neighbor embedding, we used the first 14 principal component analysis dimensions with the default settings using the RunTSNE function. For clustering, we used a resolution of 0.7 with default settings using the FindClusters function and identified tissue-specific markers expressed in the clusters using the FindAllMarkers function with default settings. The scRNA-seq data of the human reprogramming process (Liu et al. 2020) was reanalyzed under the conditions described in the paper using Seurat. Cultured monkey blastocysts (Yang et al. 2021) were reanalyzed using interactive online tools.

### WGBS processing

gDNA was extracted using DNeasy Blood & Tissue Kit, according to the manufacturer’s instructions. Library preparation and sequencing were performed by Macrogen Japan Corp. Briefly, bisulfite conversion was performed using EZ DNA Methylation-Gold Kit (ZYMO RESEARCH), according to the manufacturer’s instructions. The eluted DNA was processed using Accel-NGS Methyl-Seq DNA Library Kit (IDT), following the manufacturer’s recommendations. The libraries were sequenced on Illumina NovaSeq 6000, generating 150- base paired-end reads. The WGBS samples had a ≥99.5% sodium bisulfite conversion rate.

### WGBS data analysis

Raw sequencing reads were trimmed using Trim Galore with the recommended conditions for Accel-NGS Methyl-Seq DNA Library Kit and aligned to the human reference genome (GRCh38/hg38) using Bismark with default parameters. Bismark eliminated duplicated reads, and methylation calls were made using the Bismark methylation extractor with a minimum sequencing depth = 4 at the CpG site. CpG coverage against GRCh38 was 88% for T-iPSC^1T47^ and 87% for C-iPSC^6C6^. DMRs between T-iPSC^1T47^ and C-iPSC^6C6^ were identified using the R package DSS (≥10% difference of CpG methylation, p.threshold = 0.02, minCG = 15, and minlen = 300). The averaged methylation levels of CpGs in each enhancer were calculated using bedtools. Hypermethylated poised enhancers in C-iPSC^6C6^ compared with fibroblasts were identified using the following criteria: ≥10% higher CpG methylation in C-iPSC^6C6^ and <5% higher CpG methylation in T-iPSC^1T47^, both compared with fibroblasts. We only considered autosomal chromosomes for DNA methylation analysis.

### Characterization of the DMRs

BED files for the promoter-like signature and enhancer-like signature were downloaded from SCREEN (ENCODE3) (Consortium et al. 2020). Promoter-associated DMRs and enhancer-associated DMRs were identified by intersecting with promoter-like signature and enhancer-like signature using bedtools, respectively. DMRs within the gene bodies were identified by intersecting nonpromoter-associated DMRs with GENCODE V45 genes. DMRs overlapping the histone modifications in hESC H1 were identified using bedtools and deeptools.

### GO analysis of DEGs and DMR neighboring genes

DEGs were analyzed using Metascape for GO analysis. GREAT was used to find the genes near DMRs and to perform GO analysis. GREAT was run under the “single nearest gene” association strategy. The DMRs that associated with DEGs were selected based on the genes nearest to the DMRs, and DEGs were identified between C-iPSC^6C6^ and T-iPSC^1T47^.

### Statistical analysis of the characteristics of DMR

We used shuffled control bed files as a control for statistical analysis. We shuffled the location of the DMRs on the same chromosome using the “shuffle” tool from bedtools and recalculated the overlaps to promoter-like signature, enhancer-like signature, and gene bodies. Statistical significance was determined by the chi-square test. We used these bed files to write heatmaps of histone modification for comparison.

*Data availability*

All data supporting the conclusions are present in the paper or Supplemental Material. The sequencing data generated in this study are available through the DNA Data Bank of Japan, DRA013769 (scRNA-seq), DRA014828 (bulk RNA-seq), and DRA017148 (WGBS).

## Supplemental information

**This PDF file includes**

Supplemental Figures S1–S10.

## Other Supplemental Materials for this manuscript include the following

Supplemental methods and materials. Supplemental Tables S1–S6.

---

**Supplemental Figure S1.**
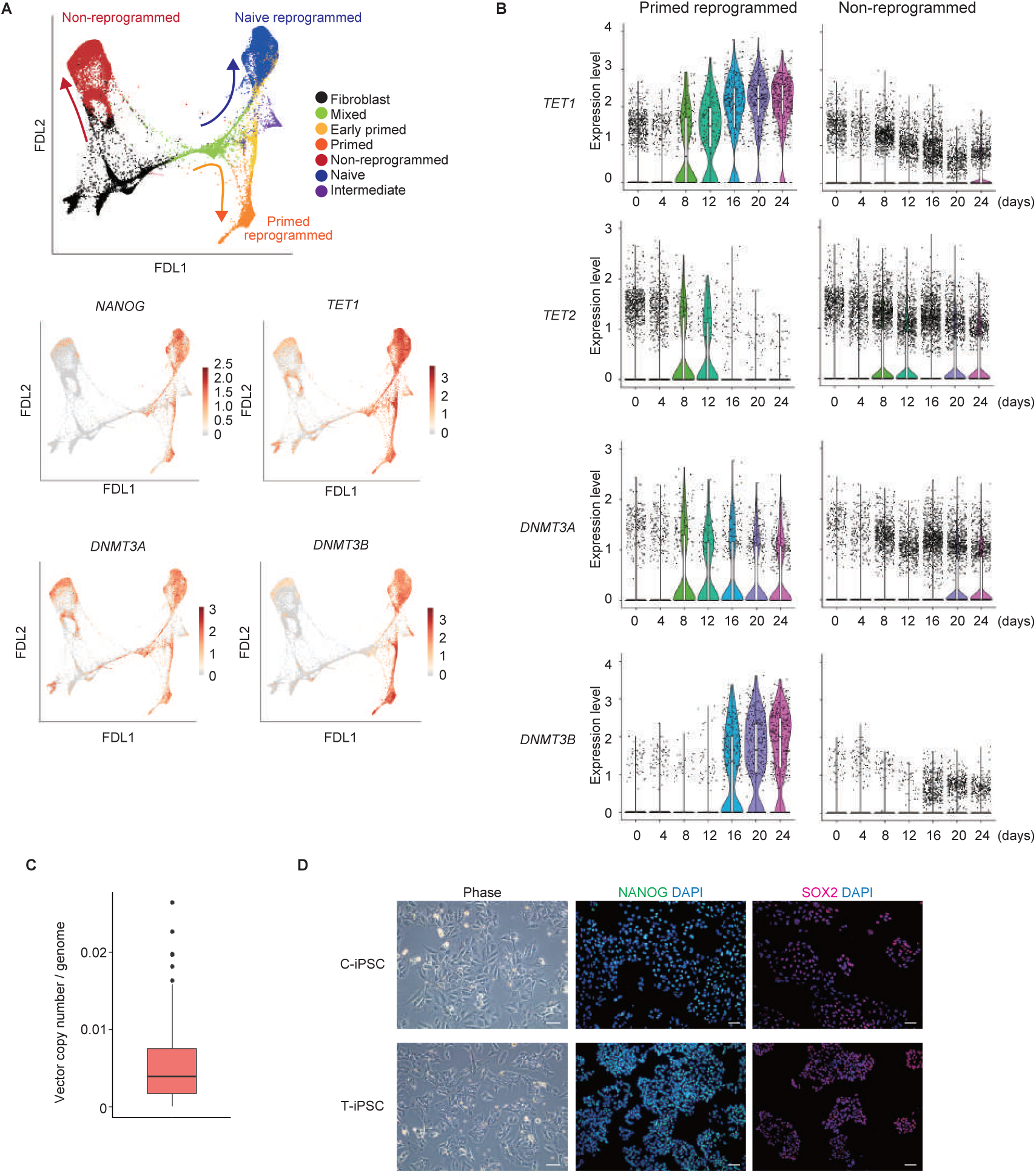
related to Figure 1. Gene expression profiles of *NANOG*, *TETs*, and *DNMT3*s during naïve and primed reprogramming from human fibroblasts to iPSCs. *(A)* Force- directed layout (FDL) visualization of the fibroblast, naïve, and primed scRNA-seq libraries (reanalyzed from Liu et al. 2020). Arrows in the upper panel show the cell trajectories to non- reprogrammed (red), naïve (blue), and primed (orange) pluripotent cell states. *(B)* Violin plots of *TET1*, *TET2*, *DNMT3A*, and *DNMT3B* expression during reprogramming. Expression data for primed reprogrammed (left) and non-reprogrammed (right) cell trajectories were scored separately. *DNMT3B*, a CpG *de novo* methylase, was induced under pluripotency acquisition (compared with the *DNMT3A* expression profile). *(C)* Box plot showing copy numbers of integrated reprogramming episomal vectors in vector-free clones determined by conventional PCR. Reanalysis using qPCR showed that conventional PCR only detected clones with >0.16 plasmid copies/genome as positive. The mean copy number of conventional PCR-deduced “vector-free” clones was 0.0062 ± 0.0067 per genome (mean ± standard deviation, n = 46). (*D)* Phase contrast images and representative immunostaining data with NANOG (green), SOX2 (red), and DAPI (blue) in established C-iPSCs and T-iPSCs. Scale bars = 50 μm.

**Supplemental Figure S2.**
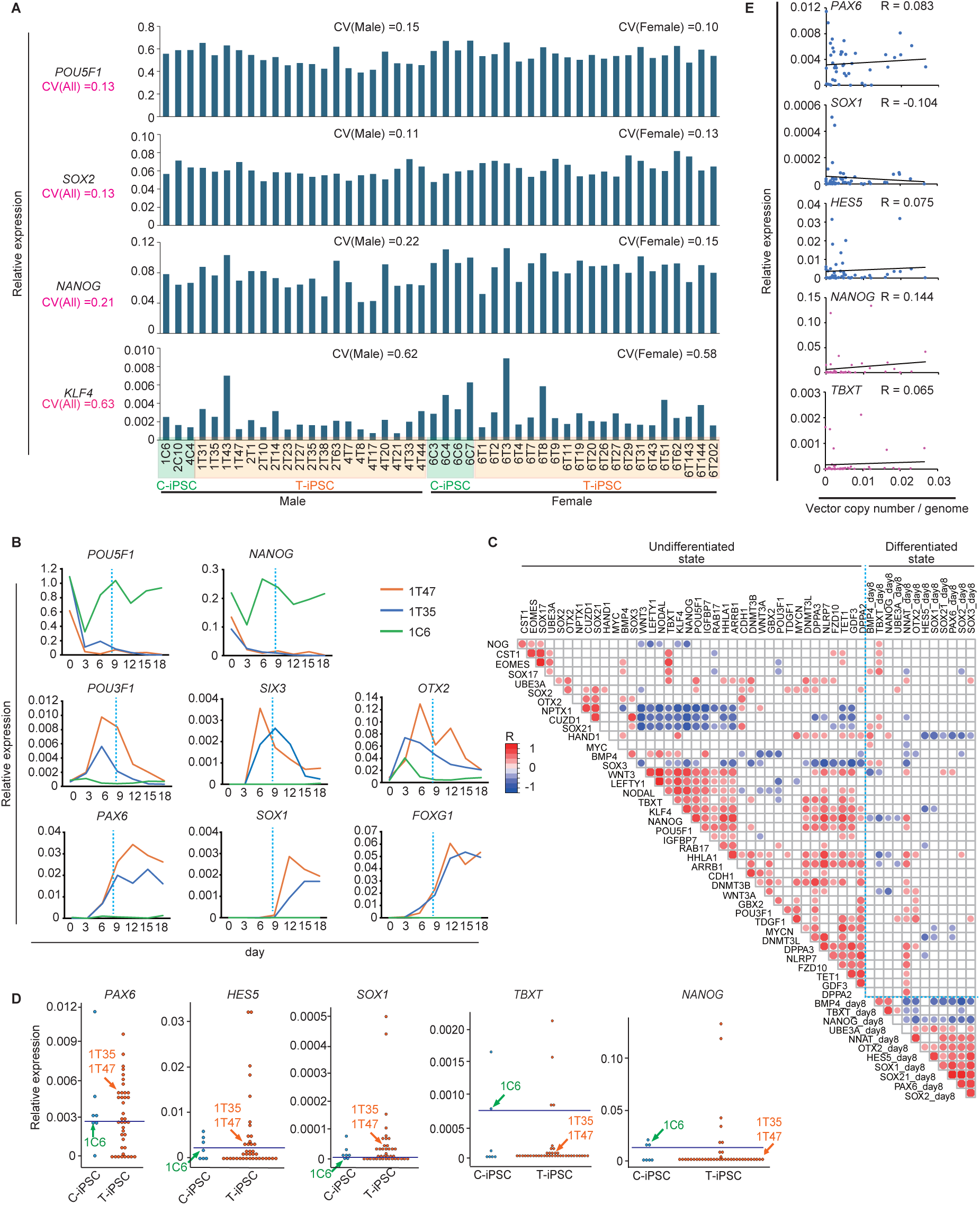
related to Figure 2. Marker profiling of each C-iPSC and T-iPSC before and after default differentiation. *(A)* Relative expression of pluripotency-associated markers on day 0 of default differentiation. *(B)* Changes in gene expression during default differentiation. The *x* axis indicates the number of days since the start of default differentiation. Day 8 of default differentiation (blue dotted lines) was the earliest reliable time to assess the differentiation capabilities of various clones (n = 1). *(C)* Correlation matrix for marker expression levels and default differentiation behaviors of all vector-free hiPSC clones. Colored circles indicate significant correlations (*P* < 0.05) between the two markers. Red and blue indicate positive and negative correlation, respectively. Color intensity and circle size indicate correlation strength (R). Blue dotted lines indicate the boundary between day 0 and day 8 expression values. *(D)* Relative expression of neural markers, *TBXT* and *NANOG*, on day 8 of default differentiation in all established clones. Each dot represents the expression level of the corresponding clone. Horizontal lines in each panel indicate the tentative threshold values for default differentiation capability. *(E)* Correlation between episomal vector copy number and neural markers *NANOG* or *TBXT* (n = 46). A residual episomal copy number of <0.03 per genome did not influence differentiation output.

**Supplemental Figure S3.**
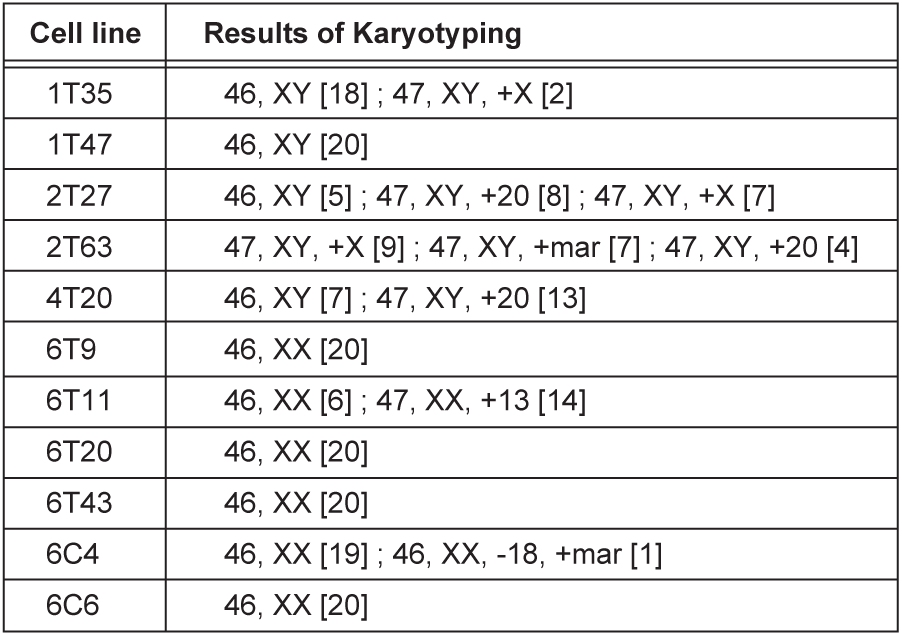
related to Figure 2. Karyotyping of default-elite clones. Numbers in square brackets represent the number of observed karyotypes in the 20 nuclei counted. +mar, marker chromosome of unknown origin.

**Supplemental Figure S4.**
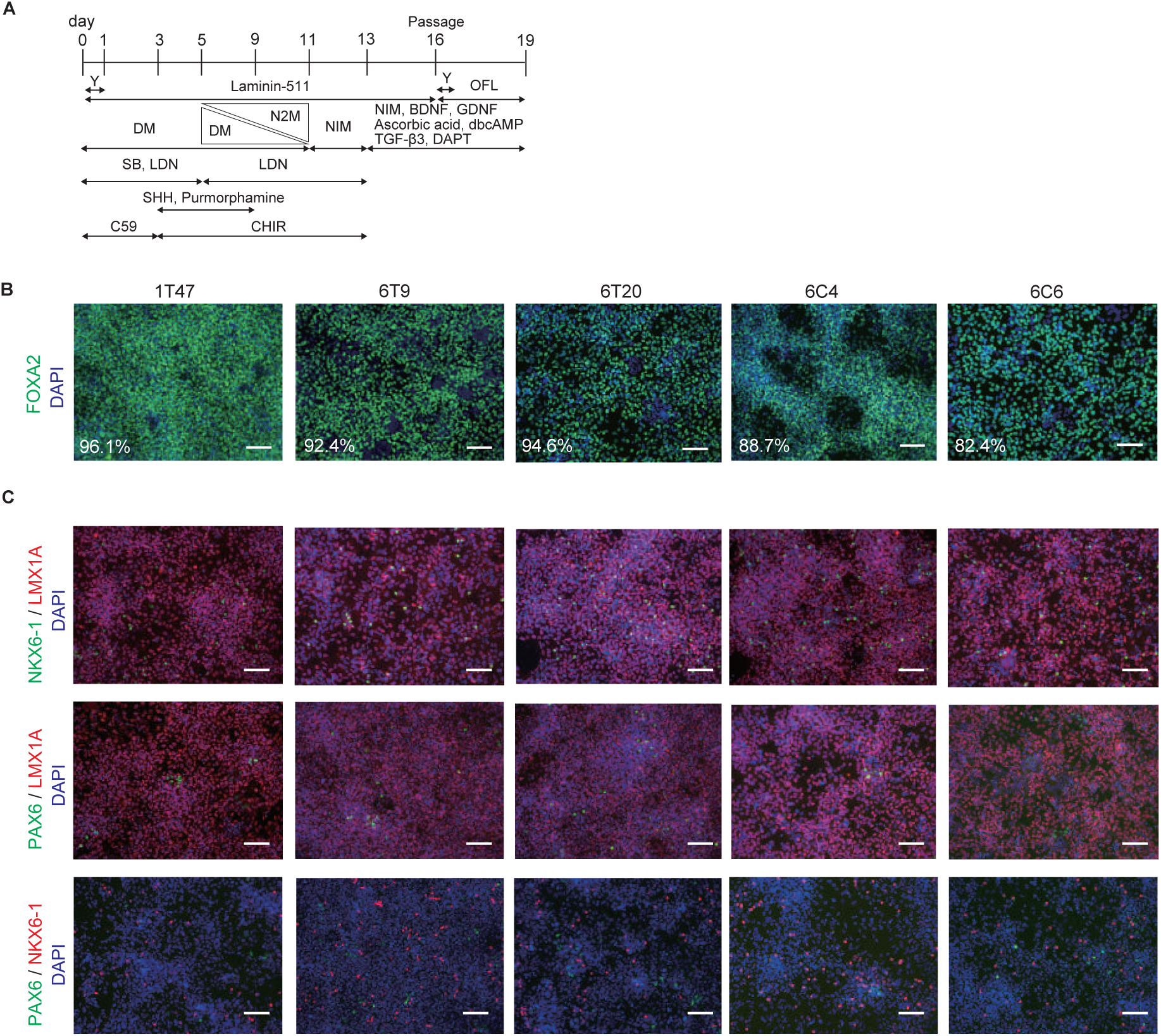
, related to Figure 3. Differentiation outputs of default-elite clones for VM progenitors. *(A)* Differentiation protocol for the induction of VM progenitors. Triple inhibition of the TGFβ-, BMP-, and WNT-signaling pathways was executed with SB, LDN, and C59, respectively. Y, Y-27632; SHH, sonic hedgehog; OFL, ornithine, fibronectin, and laminin-511. KSR-medium (DM) was gradually changed to N2-medium (N2M) from days 5 to 11, and neural induction medium (NIM) was changed from day 11. *(B)* Immunostaining images of the VM-region marker, FOXA2, in default-elite clones. Numbers indicate the percentage of positive cells. Blue, DAPI; green, FOXA2. Scale bars, 50 μm. *(C)* Immunostaining images of other VM-region markers in the default-elite clones. Blue, DAPI; green, NKX6-1 or PAX6; red, LMX1A or NKX6-1. Scale bars = 50 μm.

**Supplemental Figure S5.**
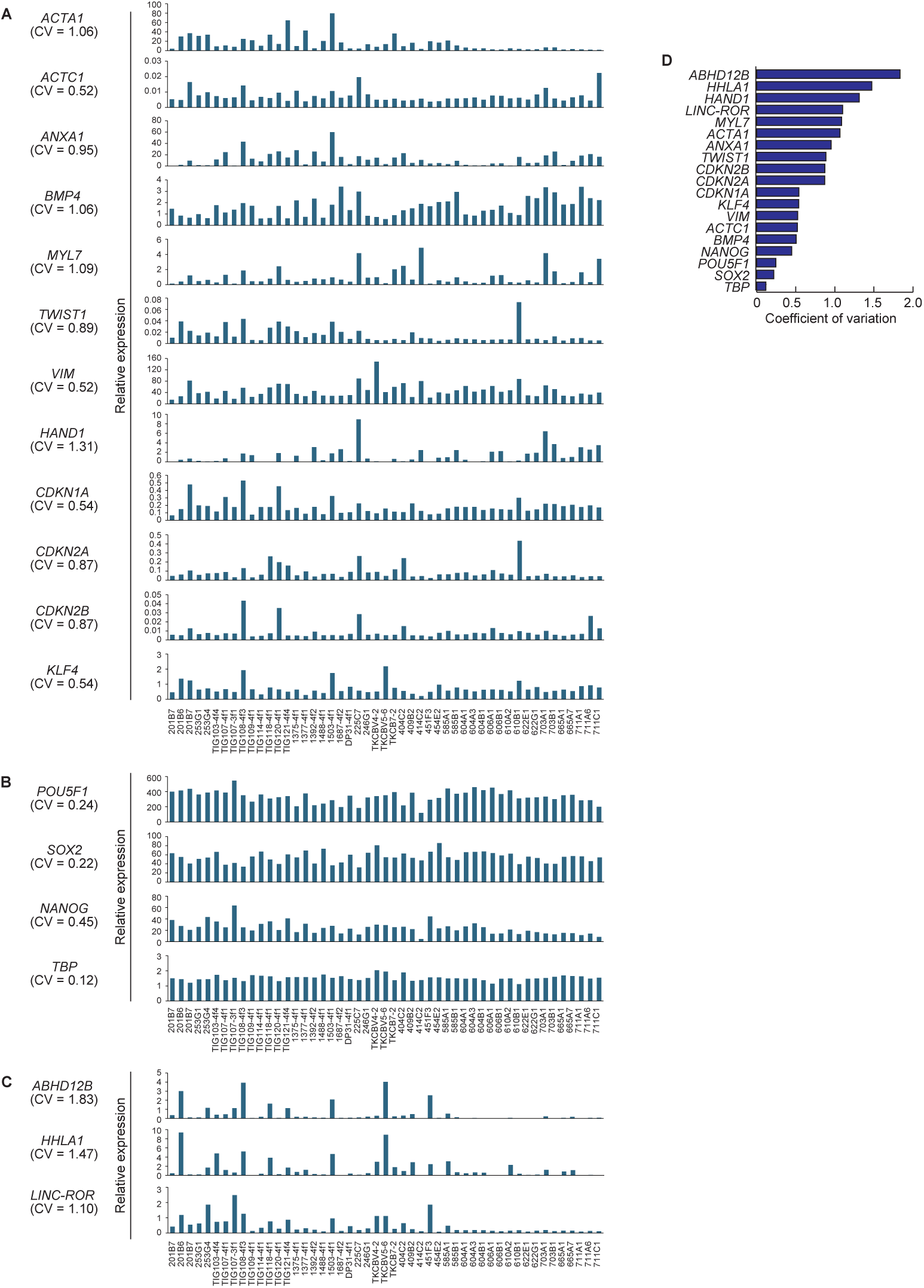
related to Figure 4. Interclonal heterogeneity of extraembryonic genes and cell cycle regulators in 49 hiPSCs (Koyanagi-Aoi et al. 2013). *(A)* Bar plots of the relative expression levels of extraembryonic genes and cell cycle regulators. *(B)* Bar plots of the relative expression levels of pluripotency-associated genes and the housekeeping gene *TBP*. *(C)* Bar plots of the relative expression levels of previously assigned differentiation-defective markers. *(D)* Bar plots of CV of gene expression for extraembryonic genes and cell cycle regulators in 49 hiPSCs. CV; coefficient of variation.

**Supplemental Figure S6.**
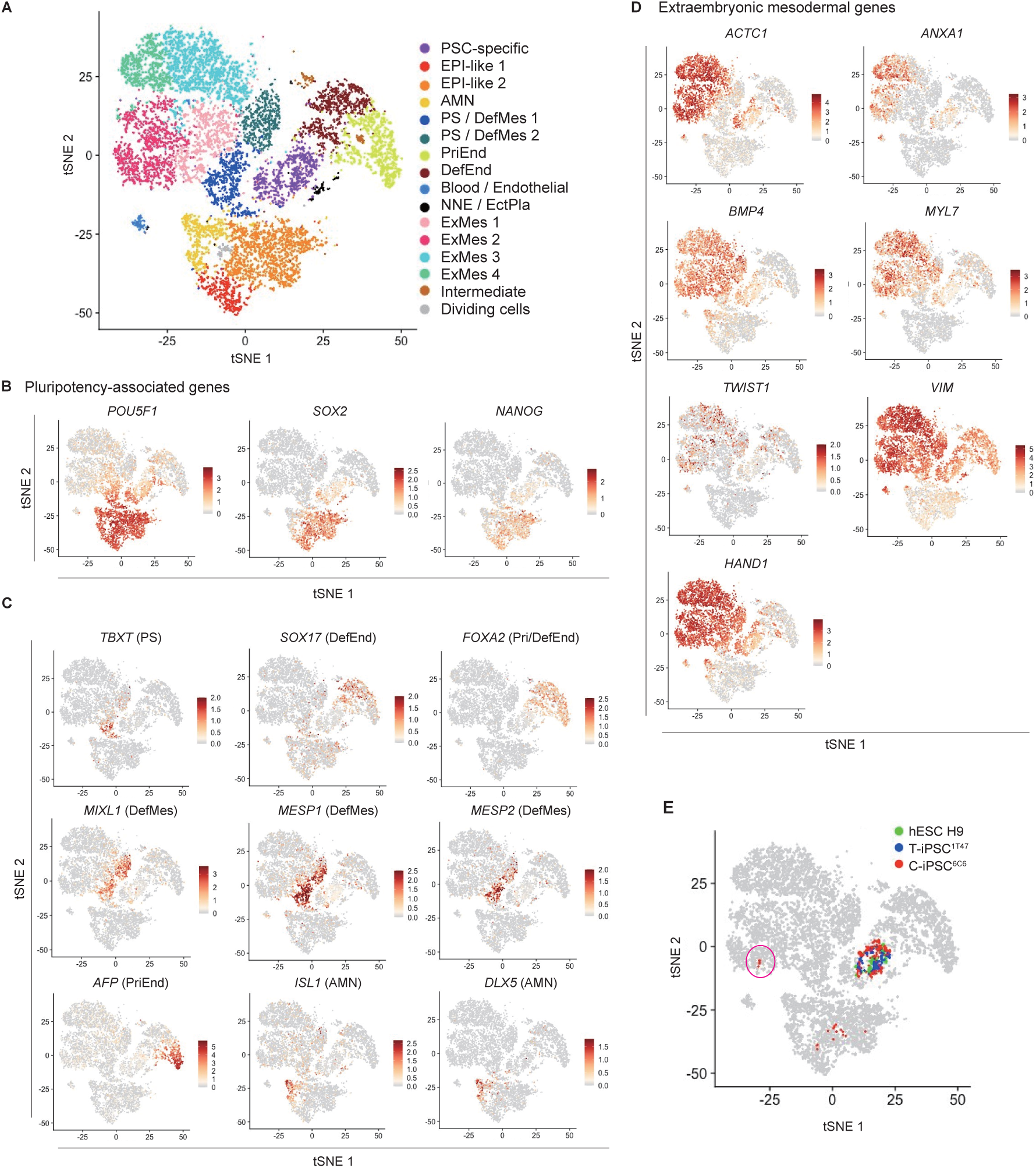
related to Figure 4. Characterization of mesenchymal genes associated with extraembryonic lineages by the reanalysis of human peri-gastruloids. *(A)* Single-cell transcriptomic integration of human peri-gastruloids (Liu et al. 2023), T-iPSC^1T47^, C-iPSC^6C6^, and hESC H9 (Messmer et al. 2019). tSNE plot of human peri-gastruloids integrated with T-iPSC^1T47^, C-iPSC^6C6^, and hESC H9. EPI, epiblast; AMN, amnion; PS, primitive streak; DefMes, definitive mesoderm; PriEnd, primitive endoderm; DefEnd, definitive endoderm; NNE, non-neural ectoderm; EctPla, ectoderm placode; ExMes, extraembryonic mesoderm. *(B)* tSNE plot showing pluripotency-associated gene expression. *(C)* tSNE plot showing the expression of *TBXT* (PS), *MESP1/2* and *MIXL1* (DefMes), *AFP* (PriEnd), *SOX17* and *FOXA2* (DefEnd), and *ISL1* and *DLX5* (AMN). *(D)* tSNE plot showing mesodermal and extraembryonic gene expression. *(E)* tSNE plot showing the locations of T-iPSC^1T47^, C-iPSC^6C6^, and hESC H9.

**Supplemental Figure S7.**
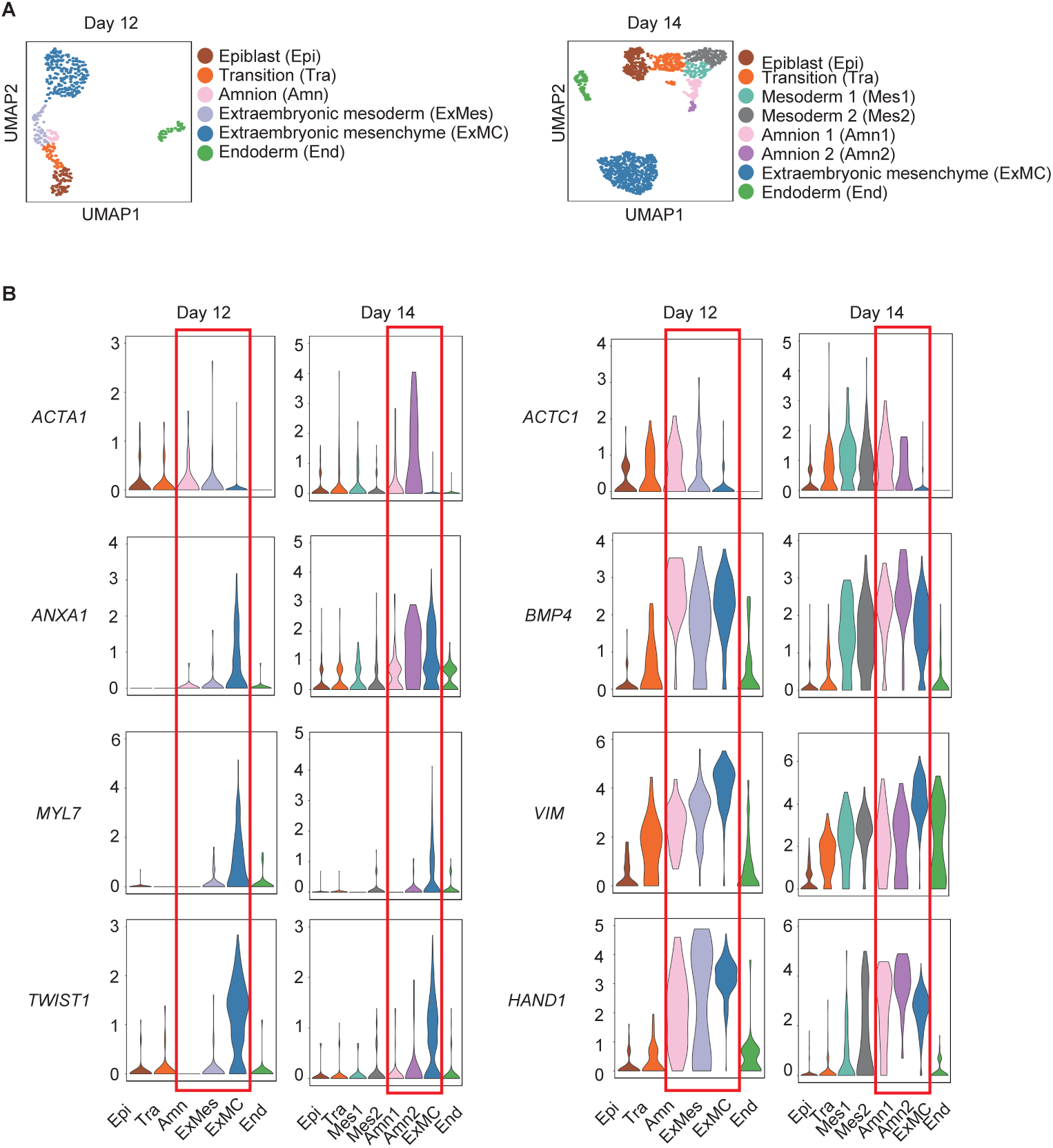
, related to Figure 4. C-iPSC^6C6^-high-DEGs are expressed in extraembryonic lineages during early monkey development (Yang et al. 2021). *(A)* UMAP plot of cells from *in vitro* cultured monkey embryos on days 12 and 14. Day-12 monkey embryos are phenotypically comparable to hPSCs. *(B)* Violin plot showing relative expression of extraembryonic-associated genes. Red rectangles surround extraembryonic lineage plots.

**Supplemental Figure S8.**
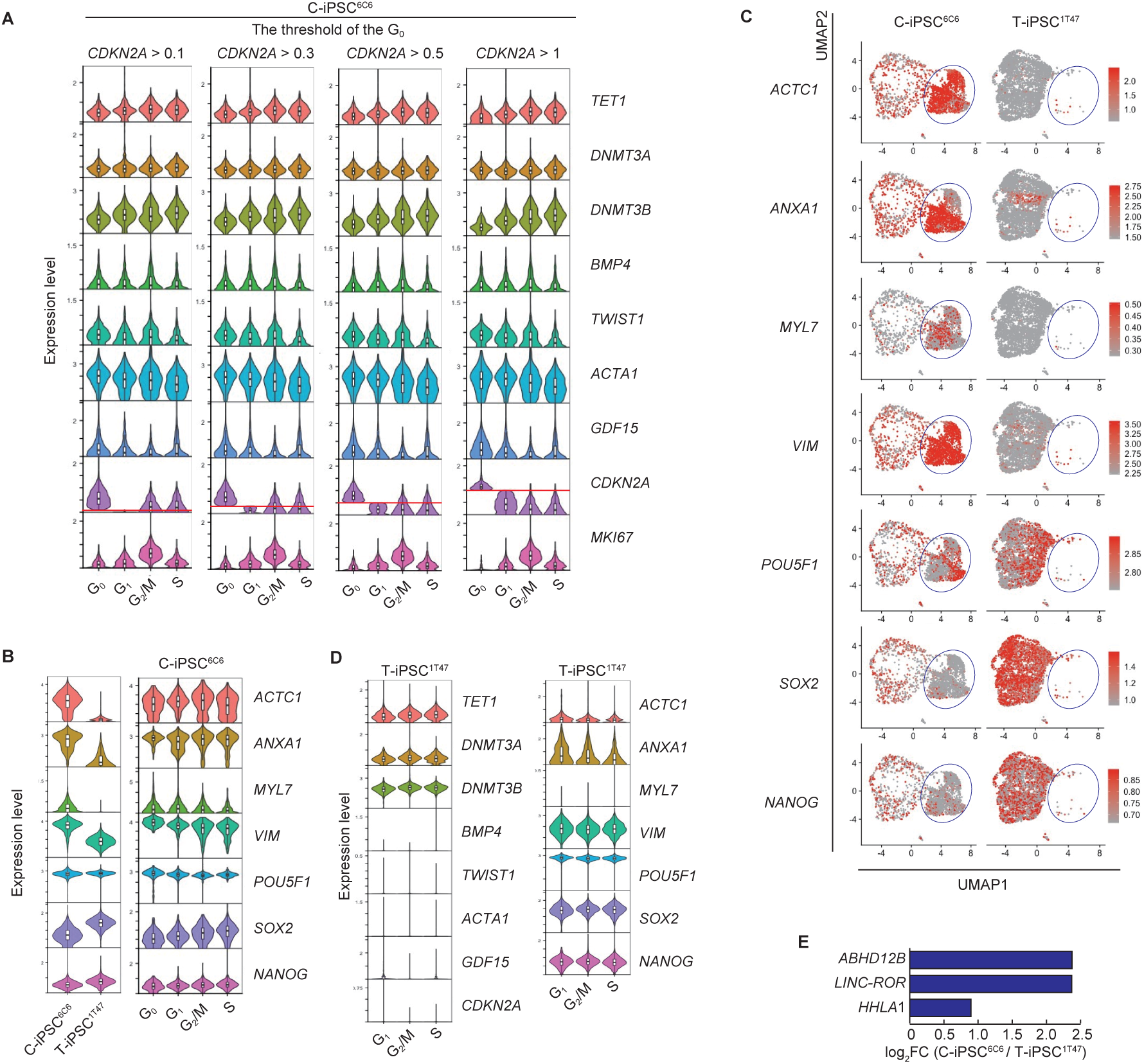
related to Figure 4. G0 phase annotation and characteristics of C- iPSC^6C6^ and T-iPSC^1T47^ in each cell cycle. *(A)* Violin plots showing the distribution of gene expression levels of regulatory genes for DNA methylation, extraembryonic, aging-, and cell cycle-associated genes in each cell cycle phase in C-iPSC^6C6^. Each panel represents differences in the distribution of gene expression for each factor when different expression levels of *CDKN2A* define the G0 phase. We defined *CDKN2A* >1 as the G0 phase. *(B)* Violin plots showing the distribution of expression of extraembryonic and pluripotency-associated genes. The left panel shows the distribution of gene expression in C-iPSC^6C6^ and T-iPSC^1T47^. Right panel shows gene expression in each cell cycle phase in C-iPSC^6C6^. *(C)* UMAP plot showing expression levels of extraembryonic- and pluripotency-associated genes in C-iPSC^6C6^ and T- iPSC^1T47^. Blue circles indicate the C-iPSC^6C6^-specific population. *(D)* Violin plots showing the distribution of regulatory genes for DNA methylation, extraembryonic, aging-, and pluripotency-associated genes in each cell cycle phase in T-iPSC^1T47^. *(E)* Bar plots of the gene expression levels of previously assigned differentiation-defective markers in C-iPSC^6C6^ compared with T-iPSC^1T47^.

**Supplemental Figure S9.**
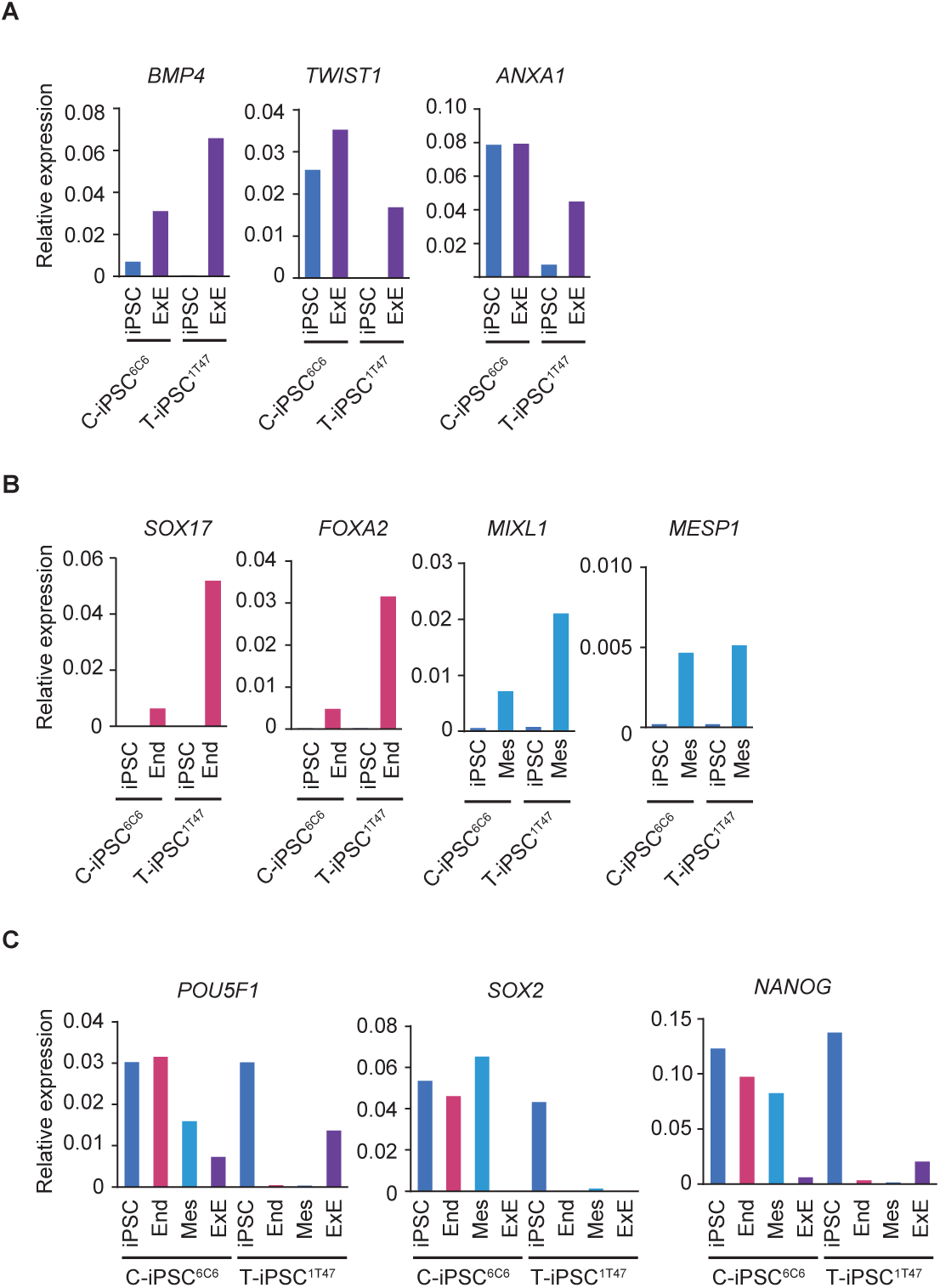
related to Figure 4. Directed differentiation into the extraembryonic and mesendodermal lineages in C-iPSC^6C6^ and T-iPSC^1T47^. *(A)* Relative expression levels of extraembryonic-associated genes in C-iPSC^6C6^ and T-iPSC^1T47^ before and after differentiation into the extraembryonic lineage on day 4 (n = 1). iPSC, before differentiation; ExE, after differentiation into the extraembryonic lineage. *(B)* Relative expression levels of marker genes after day 4 of directed differentiation into the endoderm (End) and mesoderm (Mes) (n = 1). *(C)* Relative expression levels of pluripotency-associated genes before and after differentiation into various lineages (n = 1). Pluripotency-associated markers were retained after differentiation in C-iPSC^6C6^, indicating differentiation defects.

**Supplemental Figure S10.**
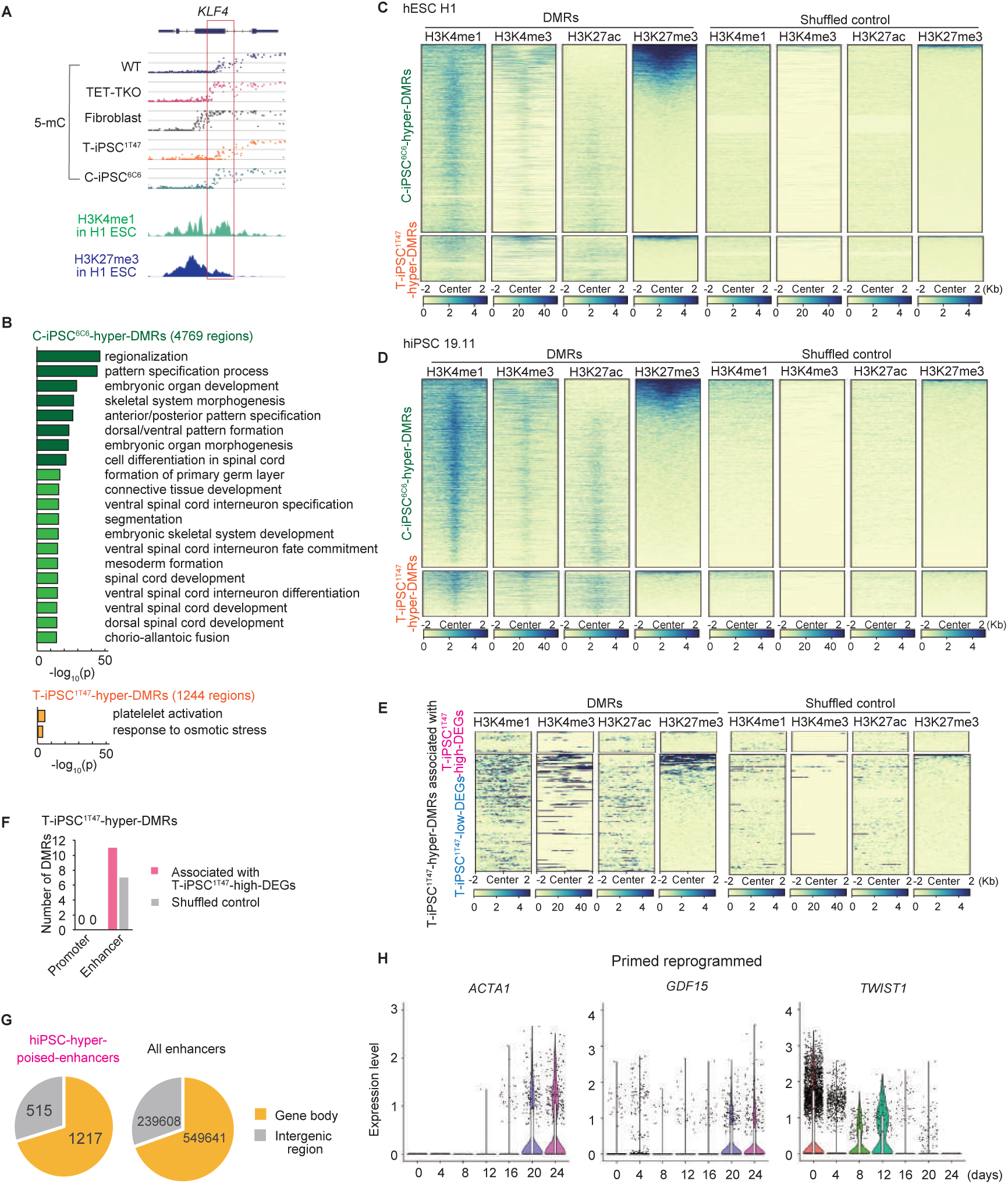
related to Figure 5. Characterization of DMRs between C-iPSC^6C6^ and T-iPSC^1T47^ and the extraembryonic-associated gene expression dynamics during reprogramming. *(A)* Genome browser tracks for WGBS data (upper panel, 5-mC) and ChIP-seq data (lower panel) of *KLF4*. Red rectangle shows the DMRs between C-iPSC^6C6^ and T- iPSC^1T47^. WT, hESC HUES8; TET-TKO, TET1/2/3-triple-knockout HUES8. *(B)* Functional enrichment of genes associated with DMRs between C-iPSC^6C6^ and T-iPSC^1T47^ using GREAT. *(C)* Heatmap of ChIP-seq signals of histone modifications (hESC H1) in C-iPSC^6C6^-hyper- DMRs and T-iPSC^1T47^-hyper-DMRs (left panel). Right panel shows histone modifications in the shuffled control regions. The rows are sorted in order of H3K27me3-densities. *(D)* Heatmap of ChIP-seq signals of histone modifications (hiPSC 19.11). *(E)* Heatmap of ChIP-seq signals of histone modifications (hESC H1) in T-iPSC^1T47^-hyper-DMRs associated with the DEGs between C-iPSC^6C6^ and T-iPSC^1T47^. *(F)* Bar plots showing the number of T-iPSC^1T47^-hyper- DMRs associated with T-iPSC^1T47^-high-DEGs overlapped in promoters and enhancers. *(G)* The proportion of *de novo* methylated poised enhancers (hiPSC-hyper-poised-enhancers) in gene bodies and intergenic regions. All enhancers described are those on autosomes. *(H)* Violin plots of *ACTA1*, *GDF15*, and *TWIST1* expression during primed reprogramming (Liu et al. 2020).

